# Coordinated gene expression variability encodes the regulatory state of cells

**DOI:** 10.1101/2025.07.09.663203

**Authors:** Fabrizio Olmeda, Yiteng Dang, Fabian Rost, Vincenzo Maria Schimmenti, Ivan Di Terlizzi, Steffen Rulands

## Abstract

Gene expression is inherently stochastic, leading to substantial cell-to-cell variability in mRNA and protein abundances. Variability in the expression of individual genes has been associated both with impaired signal processing and with facilitation of stress responses and differentiation. Here, combining machine learning, theory, and analysis of scRNA-seq data across various organisms and tissues, we show that variability in gene expression can be coordinated cell-wide. We define a statistical score that quantifies this coordination in single-cell data and demonstrate that distinct coordination patterns reflect the regulatory state of cells. We further develop a physics-informed machine-learning framework that identifies and predicts such variability patterns. Coordinated gene-expression variability emerges as a hallmark of stem and progenitor cells and distinguishes intrinsic stochasticity from cell-population heterogeneity. Together, our results establish the structure of gene expression variability as a cellular-scale signature of cell identity and regulatory organization.

## Introduction

The self-organisation of cells into complex tissues relies on the precise specification of cell states, ensuring that appropriate cellular identities are established at the correct position and time during development and regeneration [1, 2]. In multicellular organisms, cells typically follow hierarchical differentiation trajectories, progressing from stem and progenitor states toward increasingly specialised phenotypes. At the molecular level, these states are regulated by intricate gene regulatory networks (GRNs), involving transcription factor, DNA interactions, and epigenetic modifications. Therefore, gene expression is widely considered to be a primary determinant of cellular identity and function [3, 4].

Gene expression, however, is intrinsically stochastic [5], leading to substantial cell-to-cell variability. Noise at the single-gene level has been extensively studied and characterized both experimentally and theoretically [6, 7]. Recently, single-cell RNA sequencing technologies are being used to access cell-to-cell variability for a large number of genes [8]. A major challenge with these technologies is that the observed variability reflects both biological and substantial technical noise [9, 10], which requires theoretical frameworks capable of disentangling these contributions [11].

High variability of gene expression is often detrimental for biological function, an example being the precision of signal transduction or protein-protein interaction, or gene regulation [12]. Gene expression noise can, however, in itself have a biological function. Stress response genes have been shown to exhibit increased noise levels in microbes and have been suggested to support the regulation of stem cell differentiation [13, 14]. Experimental work has also shown that gene expression noise can propagate across genes [15], such that fluctuations in one gene can induce fluctuations in another gene. This raises the question of whether cell-wide propagation of gene expression fluctuations gives rise to emergent regulatory structures beyond what is captured by mean-expression or diversity-based metrics [16, 17, 18].

Here we develop a theoretical and statistical framework (Fig. 1a) validated against several scRNA-seq data sets, to show that gene–gene variability can be coordinated on the cellular scale. We define a measure (λ_max_) that quantifies the degree of cell-wide coordination in gene expression noise and investigate its association with cellular potency and different states, including dedifferentiation and proliferation, This provides a framework to distinguish whether observed variability in scRNA-seq data is due to stochastic gene expression or reflects cell-population heterogeneity [18]. We leverage annotated populations from published scRNA-seq datasets and demonstrate latent features of cellular potency that are not captured by conventional analyses. We also predict that correlations among cells within the same annotated state reflect the dynamical organisation of regulatory networks. We further use our framework to train a machine-learning model enabling data-driven inference of cell-wide properties of coordinated gene expression fluctuations [19, 20]. We apply this approach to an atlas of human cells from the gut and find that most differentiated cells exhibit a tightly constrained coordination of fluctuations, while a dedifferentiating cell population exhibit cell-wide coordinated fluctuations. This finding suggests that coordinated gene expression fluctuations capture the latent potential of cells to dedifferentiate. We finally apply our framework to scRNA-seq datasets of zebrafish heart regeneration and human pulmonary disease. We observe that injury and disease-associated cell states can be detected from variability of gene expression alone, without prior knowledge of canonical markers. Together, our findings establish coordinated gene-expression variability as a layer of regulatory organization and provide a framework to infer gene regulatory states from single-cell sequencing data, reframing how variability is interpreted in large-scale data set.

**Figure 1.**
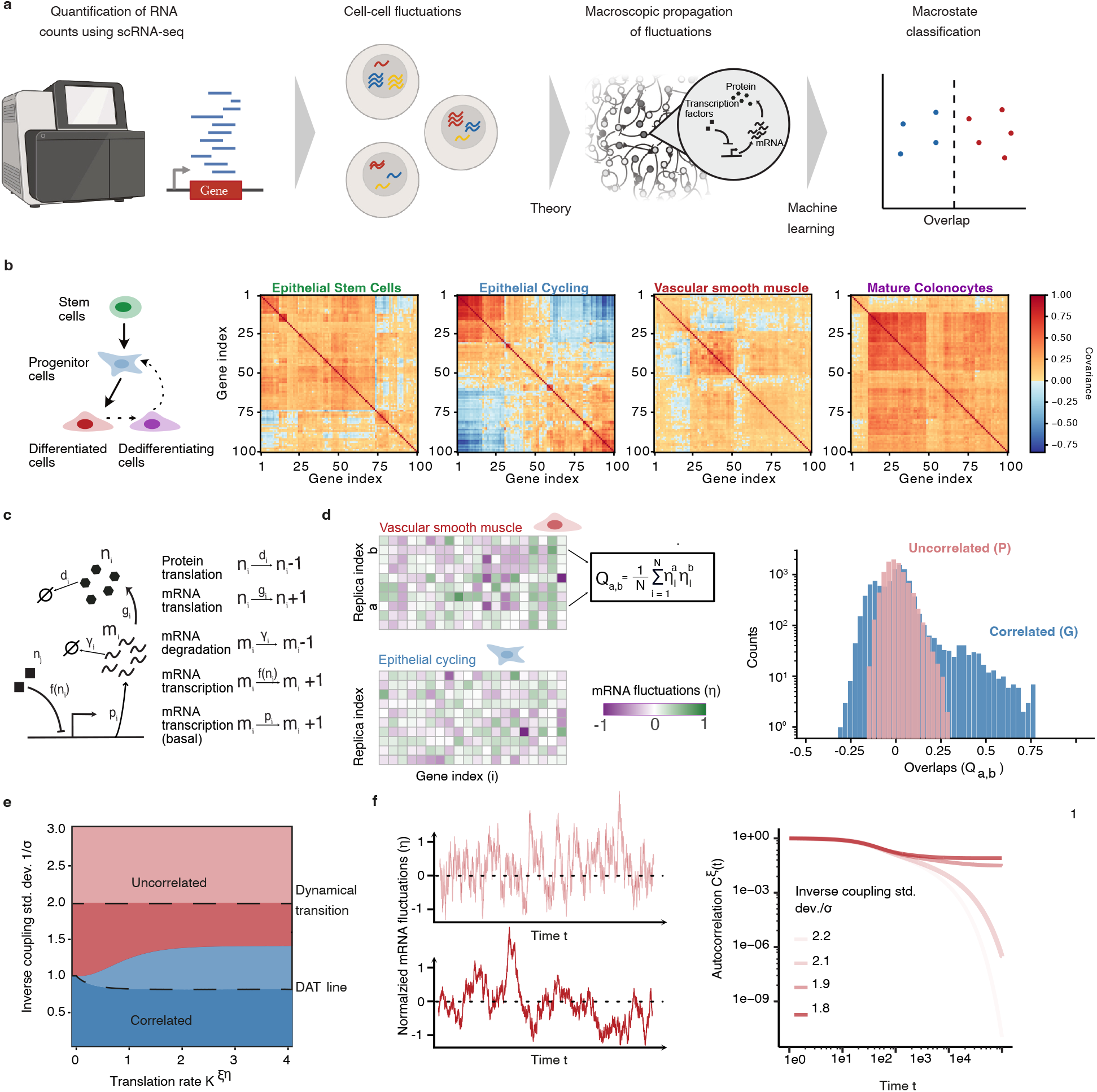
(**a**) Graphical outline of the approach used in this manuscript. mRNA at single-cell resolution is measured in RNA sequencing experiments from which we extract gene expression fluctuations. A theory for macroscopic fluctuations of gene expression allows us, using restricted Boltzmann machines, to classify cells based on the macroscopic properties of these fluctuations. (**b**) Left: Simplified schematic of a typical lineage hierarchy. Right: Typical covariance matrices of gene expression fluctuations obtained from a sequencing experiment of representative cell types of different stages of the lineage hierarchy. The structure of the covariance matrices reveals different structures and patterns which emerges with increasing cell potency (from left to right: stem cells, progenitor cells, differentiated and dedifferentiating cells from the gut [21]). (**c**) Graphical representation of the different regulatory interactions in Eq. (S.1). (**d**) Left: Two different replicas for GRNs and possible distributions of the overlaps *Q*_*a,b*_ between pairs of cells sharing the same type. Right: The width of probability density functions of overlaps reflects the phase behaviour of fluctuations. (**e**) Phase boundary between a correlated like phase (P) and a correlated phase (G) obtained theoretically in *Supplementary Information* D and *Supplementary Information* F. The upper black dashed line denotes the dynamical phase transition below which time autocorrelation functions of protein and mRNA fluctuations have a non-vanishing plateau. The lower black dashed line (DAT) denote the phase transition between correlated and uncorrelated fluctuations for the our model choice of rescaled fluctuations. The light-blue region denotes the DAT line for the choice of binary fluctuations. (**f**) Left: Typical mRNA fluctuations of a single genes over time for gene networks in the uncorrelated phase above and below the dynamical transitions. Right: Autocorrelation functions of mRNA fluctuations above and below the dynamical transition obtained from numerical simulations. Details for the processing of the data analysed in this figure are given in Methods §2.

## Results

To understand whether coordinated cell-wide variability in gene expression reflects functionally different cell states, we began by examining correlations between gene expression fluctuations from a large atlas of single-cell RNA sequencing data from the gastrointestinal tract [21]. To this end, we defined fluctuations as the scaled values of the transcripts in each annotated cell state and computed covariance matrices [22, 23], which quantify how gene expression co-varies across genes and cells; see Methods §1.1 for a more detailed discussion about technical noise and preprocessing.

The structure of these matrices reveals strong correlations in stem cells, weaker correlations in differentiated cells (Fig. 1b), and a recovery of stem-like correlation patterns in dedifferentiating cells, such as mature colonocytes. This suggests that cell states differ not only in their mean expression profiles but also in how gene-expression fluctuations are coordinated across the transcriptome. To interpret these observations, we developed a theoretical framework that connects patterns of gene expression fluctuations to a minimal model of gene regulation, enabling a quantitative interpretation of macroscopic variability in terms of collective dynamics. Conceptually, we treat cells within the same annotated regulatory state, as approximate realizations (replicas) of a shared underlying regulatory landscape. Within this widely used approximation [24, 25], cells within the same annotated state are interpreted as sampling a common underlying regulatory landscape, such that stochastic gene-expression noise generates distinct expression configurations around a shared regulatory state. Within this picture, stem-like states correspond to regimes in which fluctuations are highly coordinated across many genes, consistent with a more permissive regulatory configuration, whereas differentiation is associated with a transition toward more constrained states in which fluctuations become weaker and effectively more independent.

### Coordination of gene-expression variability

To understand how gene expression variability can coordinate on the cellular scale, as a first step, we defined a minimal stochastic description of how fluctuations around average values of gene expression propagate through gene networks. To this end, we consider a set of *N* genes. In our model, for a given gene, mRNA molecules are translated into proteins. Both proteins and mRNA molecules degrade with constant rates, Fig. 1c. Genes interact via the expression of transcription factors; hence the transcription rate of a given gene *i* depends on abundance of proteins of other genes *j* through a sigmoidal function, which summarises the kinetics of stochastic switching of promoter activation through cooperative binding of transcription factors [26]. The stochastic time evolution of the gene network is then described by the probability of finding a given combination of protein and mRNA abundances for all genes at a given point in time (*Supplementary Information*). Absolute values of mRNA and protein abundance typically depend on factors that are not considered in this model, such as epigenetic modifications of DNA and chromatin. They are also highly specific to a given cell state and they cannot be reliably predicted with the simple model defined here. However, how fluctuations in gene expression around their average values translate between pairs of genes and whether this leads to cell-wide patterns of gene expression variability is independent of these details and may lead to a robust characterization of cell states.

To demonstrate this, we calculated the predicted steady-state distribution of gene expression fluctuations around a given attractor of the gene network (*Supplementary Information A*), which in our framework defines a cell state. Specifically, the joint probability distribution of fluctuations of mRNA and protein fluctuations, *ξ*_*i*_ and *η*_*i*_, respectively, takes an exponential form, *P* (*ξ*, ***η***) ∼exp(−ℋ), with an argument given by

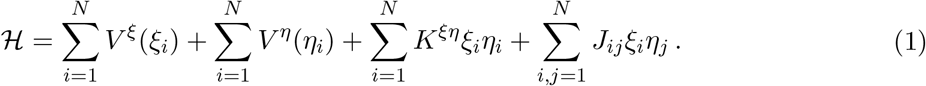

The first two terms describe intrinsic mRNA and protein fluctuations, the third term accounts for translation, and the fourth quantifies gene–gene interactions. *J*_*ij*_ encodes the effective interaction strengths between gene pairs. For the purpose of deriving testable predictions, we set *J*_*ij*_ to be normally distributed with mean zero and standard deviation 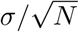 (See *Supplementary Information* B and C for a discussion of model assumptions and parameter definitions). Equation (1) has the form of the energy of a bipartite spin-glass model [27, 28]. Cellular-scale gene expression variability can be characterised in two complementary ways: gene–gene correlations, measurable from static scRNA-seq data, and the temporal persistence of fluctuations, which our framework predicts as an emergent property of the underlying network dynamics. These properties are encoded in a phase diagram.

To quantify whether fluctuations in gene expression become coordinated on the cellular scale, we first define a parameter that quantifies the degree to which gene expression fluctuations are predetermined by the interactions between genes. This parameter is given by the average correlation of fluctuations between different replicas *a* and *b*, 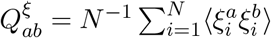 and 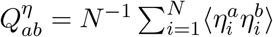, termed overlaps Fig. 1d. Biologically, overlaps measure how consistent gene expression fluctuations are between cells that rest in the same state. How overlaps change with the number of genes informs about how gene expression fluctuations propagate at the cellular level: if the overlap distribution has mean zero and standard deviation scaling as *N*^−1/2^, fluctuations are uncorrelated (*Supplementary Information* C). By contrast, a deviation from this scaling indicates a broad distribution of overlaps and the emergence of cell-wide gene-gene correlations. The boundary between these regimes (blue to red in Fig. 1e) defines a transition between phases (*Supplementary Information* D),in a model where fluctuations takes discrete binary value (Boolean). For gene-expression fluctuations, as considered here, this static transition is shifted to lower values, as indicated by the lower dashed line (DAT).

We then asked whether the dynamical properties of gene-expression variability could be inferred from gene–gene variability alone. We therefore computed autocorrelation functions of mRNA and protein fluctuations [29, 30] (*Supplementary Information* E, F). In the uncorrelated regime, autocorrelations decay exponentially with timescales set by gene-expression kinetics. If the scaled interaction strength exceeds a critical threshold *σ* > 1/2, the system undergoes a dynamical transition (Fig. 1e), marked by a persistent plateau in the autocorrelation function, which indicates that fluctuations in gene expression are long-lived with memory of expression states. The slow relaxation of fluctuations in gene expression could explain the slow differentiation dynamics of stem cells observed in [18, 31].

Numerical simulations confirm this transition and the violation of the *N*^−1/2^ scaling of overlaps (Fig. 1f). Together, these results connect overlap statistics and gene-expression variability to dynamical regimes of gene regulatory networks. As overlaps can be estimated from scRNA-seq data, our framework suggests that aspects of cellular-scale fluctuation dynamics may be inferred from static measurements and that gene-gene interactions could contribute to extended expression memory.

### Scaling analysis of gene-expression variability distinguishes stem and differentiated cells across brain development and adulthood

Our minimal model makes simplifying assumptions about the regulation of gene expression. To test its theoretical predictions empirically, we analysed publicly available single-cell RNA sequencing data from multiple tissues. However, extracting gene-expression fluctuations from such data is challenging due to substantial technical noise and biases [32, 8]. To address this problem, we employed complementary strategies designed to be robust to technical artefacts. First, we quantified how the standard deviation of the overlap distribution scales with the number of genes across several cell types. Second, we exploited the formal similarity between Eq. (1) and the energy function of a restricted Boltzmann machine (RBM) [33] to infer the covariance structure of gene-expression fluctuations. Third, we analysed the spectrum of these covariance matrices, using their leading eigenvalues as a model-free measure of collective gene-expression modes.

A unique prediction of our model is the potential emergence of cell-wide correlated fluctuations in gene expression. We therefore examined whether the variance of the overlap distribution, 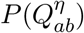, deviates from the scaling expected for uncorrelated fluctuations. Because absolute values of the variance depend strongly on technical parameters, we focused instead on scaling exponents, which are dimensionless and therefore provide robust measures of phase behaviour [34]. If gene-expression fluctuations are uncorrelated, the central limit theorem predicts that the standard deviation of the overlap distribution scales with the number of genes as *N*^−1/2^. By contrast, in a correlated phase, the decay is predicted to be slower. To test this scaling empirically, we analysed scRNA-seq data from developing and adult mouse brain [35] (Fig. 2a). For cell states that are longer lived than typical mRNA and protein lifetimes, we identified different steady states of the gene regulatory network using clustering (Fig. 2b) and computed gene-expression fluctuations within each cluster (Methods §1.1). For several cell types, the standard deviation of the overlap distribution decreased more slowly than predicted by the central limit theorem, particularly in progenitor populations such as nIPC (Fig. 2c and Fig. S1a,b). To exclude that the observed scaling directly reflects collective interactions between genes and not technical artifacts, we performed two orthogonal controls. First, if the definition of a steady state were too broad, the distribution of overlaps could be artificially wider, potentially leading to deviations from *N*^−1/2^ scaling. To test this possibility, we defined progressively smaller local neighbourhoods in gene-expression space as steady states. Deviations from *N*^−1/2^ scaling persisted across subsamples (Fig. 2d and Fig. S2a-c), indicating that the observed effect is not driven by cluster size or annotation. Second, to exclude that the deviations reflect generic features of technical noise in single-cell RNA sequencing data, we simulated single-cell RNA-seq reads assuming uncorrelated gene expression while reproducing realistic biological and technical noise (Methods §1.1 and Fig. S3a,b). These simulated data strictly followed the *N*^−1/2^ scaling predicted by the central limit theorem (Fig. S3c,d). Furthermore, to exclude that deviations reflect non-Gaussian fluctuation distributions rather than genuine correlations (Fig. S3e), we verified that non-Gaussianity does not differ across clusters in the simulated data.

**Figure 2.**
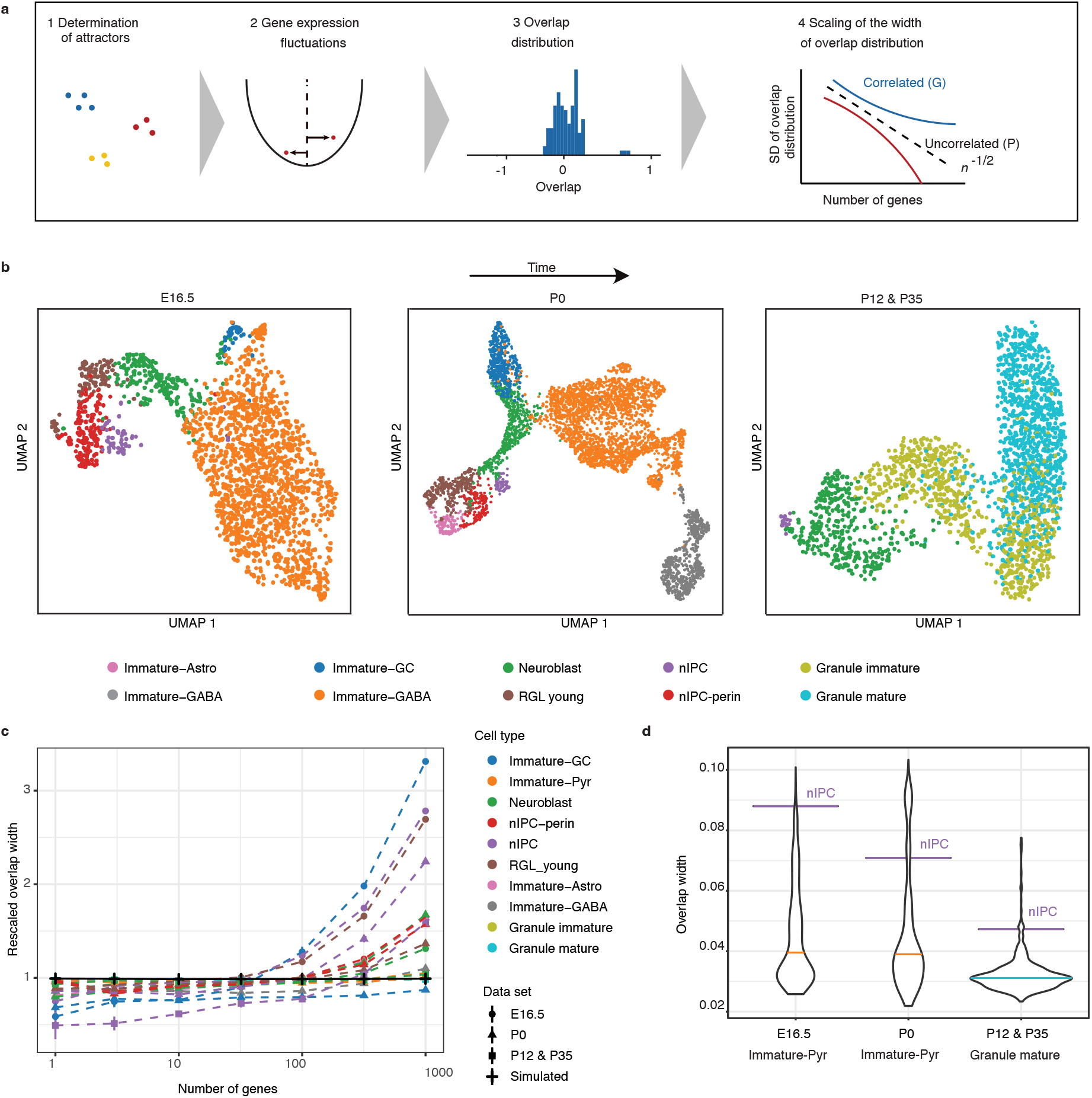
(**a**) Schematic of the computational pipeline used to detect correlated fluctuations in scRNA-seq data. After filtering out irrelevant clusters, we computed the gene fluctuations for each cell and each gene (*Supplementary Information* 1.1). From this we computed the overlap distribution for each cluster. Finally, we subsampled genes to obtain how the overlap width scales with the number of genes. The_black line shows the mean overlap width over different subsamples (blue circles). The_ dashed blue line shows the 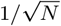 scaling obtain from the central limit theorem. (**b**) Uniform Manifold Approximation and Projection (UMAP) plot of the data at three different developmental times, with colours indicating different cell clusters. Each dot represents a cell. (**c**) The rescaled overlap width, obtained by multiplying the width of the overlap distribution by 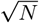, as a function of the number of genes *N*. The simulated uncorrelated data set (black line) shows a flat line, indicating that the overlap width scales as 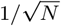 as expected. The different clusters of the single-cell data shown in (b) show different deviations from this 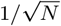 scaling. (**d**) Comparison of overlap width of the nIPC cluster to subsamples of cells from the largest clusters, Methods §1.1. The violin plots show the distributions of overlap widths obtained from 1000 independent subsamples of the clusters. Horizontal line indicates the median.

These results show that progenitor and differentiated cells differ not only in mean expression but also in the collective organization of fluctuations. Although broad overlap distributions could reflect hidden sub-structure, their persistence under subsampling argues that the observed coordination cannot be explained only by cluster heterogeneity.

### A physics-informed machine learning model infers collective gene-expression modes in the human gut

To more directly assess whether gene expression fluctuations can be coordinated on the cellular scale, we trained a physics-informed machine learning model to infer the values of the gene-gene interaction strength, *J*_*ij*_ in Eq. (1). Specifically, we noted a formal analogy between our theoretical framework and restricted Boltzmann machines (RBMs) [33]. RBMs have been applied in the context of disordered systems [36] and in biological contexts [37, 38]. An RBM is a probabilistic machinelearning model that defines a joint distribution over observed variables ***v***, and latent variables ***h***. If ***v*** and ***h*** are continuous variables, the probability distribution takes an exponential form *P* (***v, h***) = *Z*^−1^ exp(−*E*(***v, h***)), where the energy *E* is given by 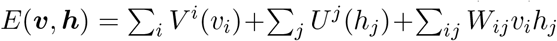, with 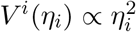and 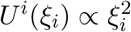, and *W*_*ij*_ the weights connecting visible and hidden units. If the number of nodes of the visible and hidden layers is identical, the RBM energy function is formally equivalent to Eq. (1). In this analogy, the observed variables correspond to mRNA fluctuations ***v*** = ***η***, and the latent variables correspond to protein fluctuations ***h*** = ***ξ***, while the weights *W*_*ij*_ correspond to the effective gene-gene interaction strengths *J*_*ij*_ in Eq. (S.5), up to diagonal terms, see Methods §2.1 for more details. Upon fitting the RBM to the data, we are therefore able to estimate the scaled standard deviation of gene–gene interactions *σ* by computing 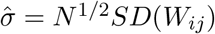, which serves as a proxy for the real *σ*. This allows us to approximately place different cell types within the phase diagram predicted by the theory (see Methods §2 for more details).

To test if cells indeed exhibit cell-wide coordination of gene expression noise using experimental data, we analysed the Pan-GI Cell Atlas [21] (Fig. 3a), which includes stem, progenitor, and differentiated cell types from the human gastrointestinal tract. We first asked if the stem and progenitor populations in the Pan-GI atlas also exhibit stronger correlated fluctuations than differentiated cells. For each annotated cell type, we computed the distribution of overlaps using an increasing number of genes (*N* = 100, 500, 1000) and quantified its width as the standard deviation of the overlaps multiplied by *N* ^1/2^. For an uncorrelated gene network, this rescaled width should not change as *N* increases. Instead, we find that it increases systematically with *N* and is consistently larger in stem and progenitor populations than in differentiated cell types (Fig. 3b), corroborating that these cells reside in a more strongly correlated regime of gene-expression fluctuations.

**Figure 3.**
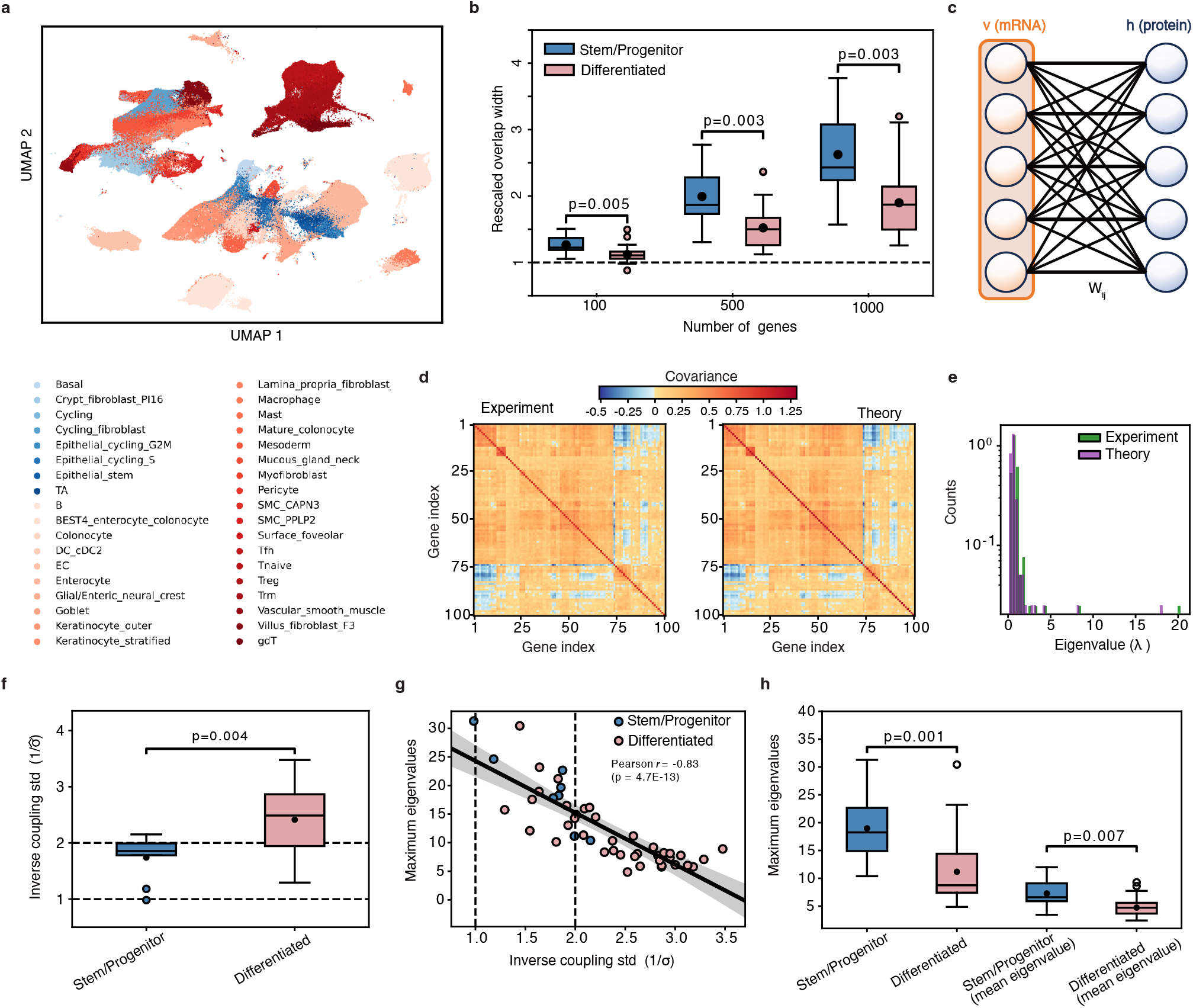
(**a**) UMAP embedding of cells from the Pan-GI Cell Atlas [21], coloured by annotated cell type; bluish hues denote stem/progenitor populations and reddish hues differentiated cell types. (**b**) Boxplots of the rescaled overlap width, defined as the standard deviation of the overlaps multiplied by *N* ^1/2^, for stem/progenitor (blue) and differentiated (red) cells, computed for *N* = 100, 500, 1000 genes. Values systematically exceed the uncorrelated expectation and increase with *N*, with stronger deviations in stem/progenitor cells. (**c**) Graphical representation of the Restricted Boltzmann Machine (RBM), where the visible layer encodes mRNA expression of individual genes and the hidden layer represents coarse-grained proteomic degrees of freedom. (**d**) Empirical gene-expression covariance matrix for epithelial stem cells compared to the RBM-predicted covariance matrix (Eq. (6)). (**e**) Eigenvalue spectra of empirical and RBM-fitted covariance matrices for epithelial stem cells. (**f**) Boxplots of the inverse coupling standard deviation 1/*σ*, inferred from the RBM, for stem/progenitor versus differentiated cell clusters; lower 1/*σ* in stem/progenitor cells indicates stronger effective couplings. P-values are from the Mann–Whitney U test. (**g**) Scatter plot of 1/*σ* versus the maximum eigenvalue of the empirical covariance matrix across cell types, revealing a strong negative correlation (Pearson *r* = 0.83, *p* = 4.7 10^−13^); the shaded band indicates the uncertainty of the linear fit. (**h**) Boxplots of the maximum covariance eigenvalue for stem/progenitor and differentiated cells. The first pair uses the 100 marker genes with highest Wilcoxon score, and the second pair reports the mean maximum eigenvalue from 100 random subsets of 100 genes drawn from the 1000-gene panels. In both cases, stem/progenitor cells show systematically larger eigenvalues (Mann-Whitney p-values indicated).

We then applied the RBM approach to the same data. For each cell type, we trained an RBM, visualised in Fig. 3c, on a selected set of 100 genes, as described in Methods §2. The model accurately captures the empirical covariance structure of gene–gene fluctuations, which is well approximated by the theoretical expression, Eq. (6) in Methods, evaluated using the fit parameters. The agreement between empirical and model covariances is shown for epithelial stem cells in Figs. 3d and S4. Covariances, interaction matrices, and code for all other clusters are available in [39].

All clusters show 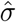 values close to one, suggesting proximity to the predicted phase transition (Fig. 1d) and motivating us to examine its biological relevance for cell identity regulation. Specifically, to test whether 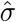 is higher in differentiated than in more potent states, as suggested by Fig. 2 and Fig. 3b, we compared 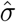between the same two cell classes. Consistent with the analysis in Fig. 3b, Fig. 3f shows that 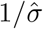 is significantly lower in stem and progenitor cells compared to differentiated ones (*p* = 4 · 10^−3^, Mann-Whitney U test), indicating greater variability of interactions and stronger effective couplings in the more potent populations.

Finally, we ask whether the value of 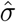 can be linked to the degree of coordination in gene-expression fluctuations among genes. Rather than analysing individual gene–gene pairs, we seek a compact descriptor that captures the emergence of dominant cell-wide patterns of gene expression variability. To this end, we considered the largest eigenvalue of the empirical covariance matrix of gene expression fluctuations, which, mathematically, measures the strength of the most prominent collective fluctuation mode. Intuitively, for a covariance matrix, one can define so-called eigen-directions which represent statistically independent modes of variation gene expression fluctuations. Eigenvalues characterise how strongly a covariance matrix changes these eigenvectors when multiplied with them. The largest eigenvalue of the covariance matrix of gene expression fluctuations, *λ*_max_, which corresponds to the first principal component, thus measures the strength of the most prominent mode of coordinated gene-expression fluctuations, providing a compact descriptor of emergent system-wide variability. In random matrices, the eigenvalues follow characteristic distributions such that values isolated from the bulk of the spectrum (Fig. 3e) signal the presence of structured correlations in the underlying data [40, 41], an approach also employed in the analysis of RBMs [42]. Across Pan-GI cell types, we found a strong negative correlation between the leading eigenvalue of the covariance matrix *λ*_max_, and the inverse standard deviation of inferred gene-gene interaction strengths, 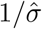 (Pearson *r* = −0.83, *p* = 4.7 · 10^−13^; Fig. 3g). This indicates stronger, more coordinated gene-expression fluctuations in highly potent cells. Because *λ*_max_ is computed directly from the empirical covariance matrix and remains well defined even for small clusters, its strong anti-correlation with RBM-based estimates supports its use as a simple proxy for collective gene-expression variability.

To test whether large eigenvalues of the covariance matrix actually probe the gene network as broadly as possible, we compared *λ*_max_ between stem/progenitor and differentiated populations using two complementary gene panels. First, we calculated *λ*_max_ for each cell type from the 100 marker genes used for the RBM analysis. Second, we repeated the analysis for 100 random subsets of 100 genes drawn from a larger set of 1000 representative marker genes and averaged the resulting values of *λ*_max_ (Fig. 3h). In both cases, stem and progenitor cells exhibit significantly greater values of *λ*_max_ than differentiated populations. This demonstrates that stronger coordination of gene expression fluctuations is a general property of the gene-regulatory networks of stem and progenitor cells, rather than an artifact of specific marker choices. Because progenitor populations often lie along continuous differentiation trajectories in transcriptional space, elevated covariance could reflect residual population heterogeneity, that is, diversity across cells in their fluctuation structure. However, its consistent enrichment across independent datasets, including stem populations, indicates that enhanced fluctuation coordination is a recurrent property of potent regulatory states rather than a geometric or dataset-specific artifact.

Despite the strong association we observed between covariance eigenvalues and cell-wide coordination of gene-expression fluctuations, we identified differentiated cell populations that nevertheless exhibit stem/progenitor-like fluctuation patterns (Fig. 3g,h). We asked whether these were merely statistical outliers, which are expected in sequencing data, or whether they encode additional biological information. To our surprise, we found that the outlier clusters correspond to colonocytes, a cell type that has been shown to retain the capacity to dedifferentiate upon injury [43]. Having validated our theoretical framework using restricted Boltzmann machines, which accurately reproduce covariance structures, we next asked whether this approach could reveal functional states within differentiated cell populations.

### Coordinated gene-expression variability in injury and disease

We then wanted to test if our framework could be further used to infer the cellular states upon injury and disease conditions. To assess this, we analysed scRNA-seq data from zebrafish heart regeneration [44], Methods §1.3. Figure 4a shows annotations reported in the original study, including dedifferentiating, proliferative, and regenerative states across multiple cell types. We asked whether an analysis based only on the covariance structure of gene expression fluctuations could recover such state changes. For each annotated cluster and time point, we computed the covariance matrix of gene expression fluctuations of marker genes and extracted its largest eigenvalue, *λ*_max_. Across lineages, *λ*_max_ peaked after injury (Fig. 4b). In Fig. S5a, we show the p-values obtained for individual clusters across the time course using an ANOVA test, with statistical significance assessed by bootstrapping over subsets of cells. Our results indicate that activation during regeneration is associated with more strongly coordinated fluctuations regime. We found that annotated subpopulations associated with distinct activation states, such as proliferative macrophages, regenerative fibroblasts, and proliferating cells, consistently displayed elevated *λ*_max_ compared to their counterparts, suggesting that *λ*_max_ could be used to distinguish activation states during regeneration.

**Figure 4.**
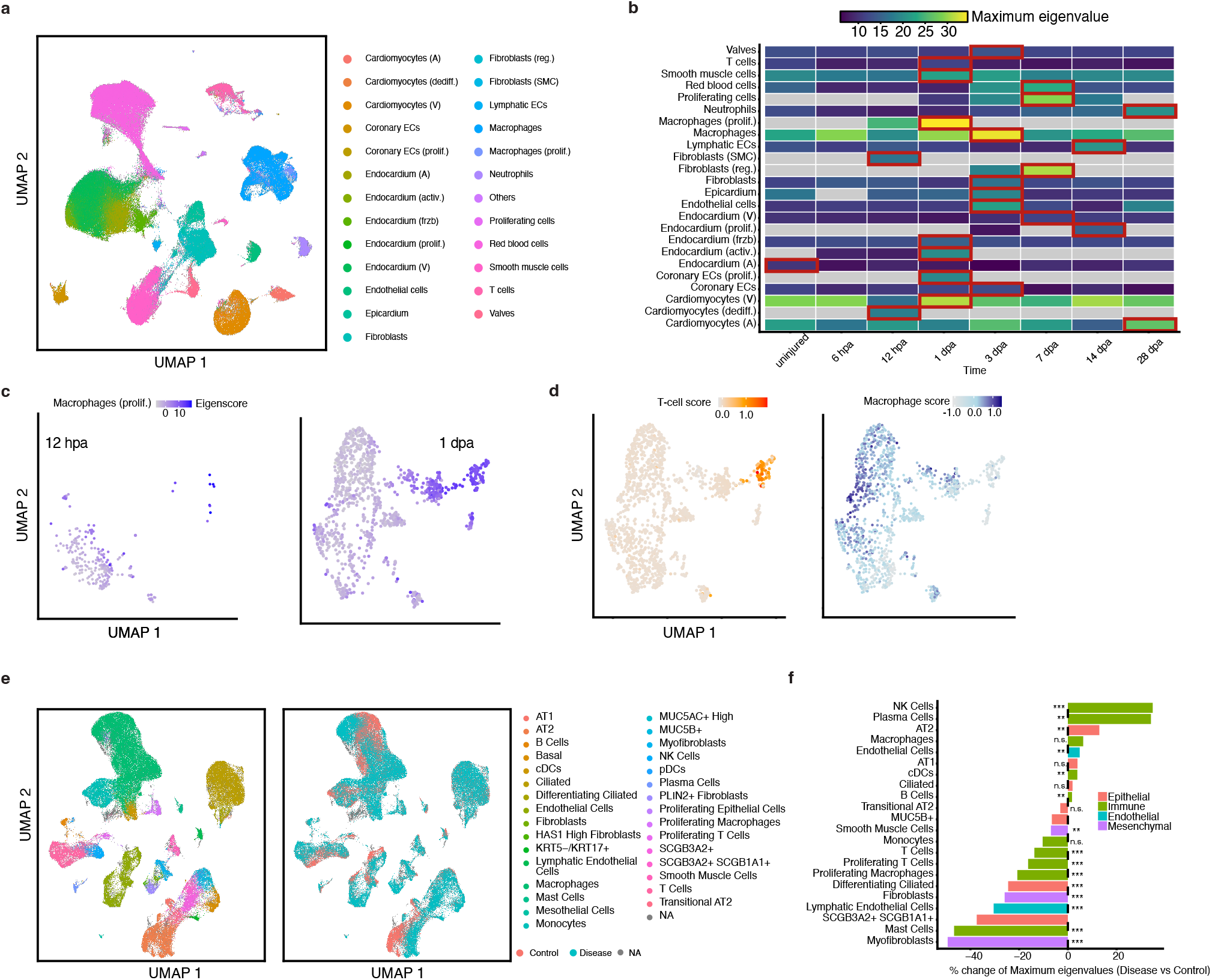
(**a**) UMAP dimensional reduction with annotated cell states, reproduce from [44], of regenerating zebrafish hearth. (**b**) Maximum eigenvalue of the covariance matrix compute for each cluster at different time points after amputation (dpa). In red we highlight the time at which the covariance matrix of each cluster is more structured, i.e. the maximum of *λ* _max_. Grey squared correspond to absence of the cluster at a given day. (**c**) Projection on the eigenvector (eigenscore) associated to the maximum eigenvalue reveals a sub-cluster of cells 1dpa in proliferating macrophages. (**d**) Scoring for T-cells and Macrophages programs reveals that this sub-cluster was caused by a T-cell population initially assigned to the Macrophage cluster (Methods §1.3). (**e**) UMAP dimensionality reduction, reproduced from [45], of scRNA-seq data from human pulmonary disease. Left: annotated cell types; Right: cells are coloured by healthy or diseased condition. (**f**) Relative change of the maximum eigenvalue for each cluster in (c) between control (left) and disease (right) conditions. Numerical p-values are shown in Fig. S5b and are calculated with bootstrapping as in Methods §1.3. Clusters lacking p-values are due to low cell numbers such that no meaningful p-values with bootstrapping methods can be obtained.

Among all clusters, the proliferating macrophages clusters has the highest value of *λ*_max_. A large value of *λ*_max_ can signify coordinated gene-expression variability as in Figs 2 and 3. Alternatively, it could reveal undiscovered substructures within the annotated cell populations. To distinguish between both scenarios, we defined an *eigenscore* for each cell. Mathematically, the *eigenscore* is obtained by projecting its gene-expression fluctuation profile onto the eigenvector associated with *λ*_max_ of the covariance matrix. Intuitively, this *eigenscore* measures how strongly a cell’s gene expression fluctuations resembles the typical fluctuation profile of the entire cluster. If a large value of *λ*_max_ for a given cell cluster were driven by a small subpopulation, *eigenscores* would segregate in transcriptome space. Instead, a homogeneous distribution of *eigenscores* indicates that coordinated variability is a global property of the cluster rather than a consequence of residual heterogeneity (Methods §1.3). We computed eigenscores for all cells and found no evidence of sub-clustering (see Fig. S5b,c for two examples), with the exception of proliferating macrophages, where the score was specifically enriched in a small subpopulation of cells (Fig. 4c). Differential expression analysis indicated strong enrichment of canonical T-cell receptor signalling genes, including Zap70, Lat, Lck, and Xd247. To determine whether this reflected heterogeneity or lineage contamination, we computed module scores for macrophage-specific (Peg1.1, Csf1ra) and T-cell-specific gene programs, Fig. 4d. Highly scored cells displayed elevated T-cell scores while lacking macrophage marker expression, and these programs were mutually exclusive at the single-cell level. These results indicate that this eigenvalue analysis can reveal hidden subpopulations or misannotation.

To further explore whether this eigenvalue analysis could enable the detection of heterogeneous sub-populations, and uncover disease-dependent differences in gene expression variability, we analysed scRNA-seq data of human pulmonary disease [45], Methods §1.3. Using the annotation of the original publication, we subdivided each cluster by condition (control vs disease), Fig. 4e, and computed gene expression variability as in Methods §1.1. We then calculated the maximum eigenvalue of the covariance matrix for each cluster. The eigenscore analysis as above did not indicate any misannotated cell clusters. We found that the maximum eigenvalues are on average greater in control compared to disease, Fig. 4f and Fig. S5c for computation of p-values. Together, these results show that our framework offers a simple, general way to detect functional state changes, such as regenerative activation or immune response. These functional state changes are encoded in collective gene-expression variability, complementing conventional transcriptomic analyses.

## Discussion

In this study, we integrate concepts from statistical physics with statistical learning and singlecell genomics to investigate how gene-expression variability relates to cellular potency. Using a minimal model of gene regulatory networks with heterogeneous and asymmetric interactions, we identify a transition between regimes of weakly and strongly correlated gene-expression variability. We show that this transition leaves robust signatures in gene-expression covariance structure that can be detected from static single-cell RNA sequencing measurements. Applying this framework to several datasets from diverse organisms and tissues, we find that stem and progenitor populations exhibit highly correlated fluctuation patterns, while differentiated cell types display less structured and weakly correlated variability. Several other methods aim to estimate cellular potency from single-cell data, including entropy-based measures [16] and transcriptional diversity metrics such as CytoTRACE [17]. These quantify expression dispersion within individual cells. In contrast, our framework measures coordinated multi-gene fluctuations across cells within a state. Thus, two populations with similar transcriptional entropy may differ in fluctuation coherence, indicating that coordinated variability provides complementary information beyond single-gene statistics.

To identify correlated fluctuations, we trained RBMs directly on scRNA-seq data. This probabilistic model mirrors our theoretical framework and learns gene co-fluctuation patterns [28]. We found that high-potency cell types, such as stem or progenitor cells, display broader coupling distributions and more structured covariance patterns, consistent with the stronger correlation regime predicted by our theory. Differentiated cells show narrower couplings and weaker co-fluctuations; interestingly, some annotated differentiated cells display stem-like fluctuation signatures.

We then ask if we could use our framework beyond categorization of cellular potency and discover functionally different cell states. To this end, we used published scRNA-seq on zebrafish hearth regeneration and human pulmonary diseases, where several groups of functionally distinct cell states were identified. We show that our framework correctly identifies cluster and sub-clusters of cell states with proliferating, dedifferentiating, and injury/disease response. In [39] we provide a function to detect covariance and eigenvalue statistics for scRNA-seq data sets.

Our results suggest that the identity of cells is defined not only by absolute concentrations of transcripts, but also by how fluctuations propagate at the scale of the nucleus. Because these fluctuations can be long-lived, they may potentially be used by cells to encode biological information [46, 47, 48]. Whether strongly correlated gene expression is indeed exploited by cells to perform biological functions is an interesting question for future research. Our work indicates a way for the biophysical interpretation of widely used single-cell sequencing experiments: although absolute gene expression values are usually strongly confounded by technical factors and highly specific to a given cell type, the way fluctuations propagate on the GRN scale gives a robust and interpretable readout of cell states. Taken together, our findings challenge the dichotomous view of stem versus differentiated states and point to a more complex organization of cell identity [49].

## Supporting information

Supplementary Figures and Theory

## Acknowledgments

F.O. received funding from the European Union’s Horizon 2020 research and innovation programme under the Marie Sklodowska-Curie grant agreement No 101034413. S.R. received funding from the European Research Council (ERC, grant agreement no. 950349. The authors thank Tuan Minh Pham for useful discussions. Figure 1a has been created with BioRender.com.

## Authors contributions

F.O. designed, planned, and conducted the research; F.O. and S.R. conceptualized and supervised the work; F.O. performed the theoretical and numerical work; F.O., Y.D. and F.R performed bionformatics and data analysis; I.D.T. designed the statistical learning pipelines; I.D.T. and V.S. performed bionformatics and machine learning; F.O, I.D.T. and S.R. wrote the manuscript; S.R. acquired funding.

## Methods

### 1 Processing of single-cell RNA sequencing data

#### 1.1 Developing mouse brain

Here we discussed the bionformatics pipeline for Fig. 2.

In order to relate our theory to experimental data from single-cell RNA sequencing experiments, we need to specify our assumptions. First, we omit the average over the disorder, meaning that every thermodynamical quantities must be computed averaging over all the possible realisation of the interaction couplings *J*_*ij*_ in Eq. (1). We have then found constraints of where observables should be computed, which means that we need to find attractors of the gene networks dynamics, which are identified as cell types. We finally have to compute overlaps for pairs of replicas, which are now identifiable as different individual cells within the same cell types, and finally get the overlap distributions.

We first utilised previously published single-cell RNA-seq data from the developing mouse brain [35] at three different embryonic stages: E16.5 (pre-natal), P0 (at birth) and P12 & P35 (post-natal). As these data sets have already been pre-processed and clustered, the only additional pre-processing steps we perform consist of filtering out genes expressed in < 20 cells and removing small clusters which are disconnected from the main cluster on the dimensionality reduced UMAP plots. This allows us to focus on clusters which are thought to be directly related to one another through cell differentiation.

We interpret each cluster as a separate stationary states and define the fluctuations and overlaps with respect to these stationary states. We first selected the 1000 most highly variable genes for each data set. We define the fluctuations by subtracting the average expression level over the cluster for each gene. Because overall gene expression levels and therefore the fluctuations can differ significantly between different genes, we also rescale the fluctuations so that they have unit standard deviation for each gene before computing the overlaps 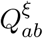. Here, the pairs of cells in the same cluster are considered to be statistically independent replicas *a* and *b*. By considering the overlaps for each pair of cells *a, b* of a given cluster, we obtain the overlap distribution of the cluster. The overlap width is then simply defined as the standard deviation of this distribution. We note that while for the theoretical constraints used before the overlap is constrained between -1 and 1, here we use the biological normalization per gene. With this choice the bound on the overlap is not strictly enforced, but does not change any result as we are interested in comparison between different cell states. Moreover, we found very few clusters which actually exceed such boundaries. In order to study how the overlap width scales with the number of genes, we subsampled genes by randomly selecting a subset of *n* < 1000 genes from the set of 1000 highly variable genes, and computed the overlaps on this subsample of genes. From this we obtained the standard deviations of the overlap distributions (overlap widths) and studied how these overlap widths scale with *n*. In order to rule out additional biological and technical effects that may lead to deviations from the central limit theorem in the overlap width scaling, we performed two additional checks. First, we noted that different clusters have different sizes (number of cells). Furthermore, the clusters that seem to show the strongest deviations from 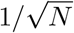 scaling seem to be predominantly the smaller clusters (Fig. 2c). We hence hypothesized that a possible confounding variable could be the size of the cluster and devised a method to control for this. Suppose we want to compare clusters of sizes *S*1 > *S*2. The idea is then to take subsamples of size *S*2 from the larger cluster with size *S*1 to match the number of cells of the smaller cluster. However, note that larger clusters could exhibit greater variability between cells even in the case where each cluster is identically distributed, simply due to the fact that taking more samples from a distribution increases the chance of hitting the tails of the distribution. Therefore, we restricted our subsamples to cells which are sufficiently similar to one another. We used the fact that in order to compute UMAPs, one first needs to compute a neighbour graph, whereby each cell is connected to k nearest neighbours (kNN). We hence took kNN neighbourhoods of size *S*2 of randomly sampled cells from the cluster of size *S*1, and computed the overlaps on the groups of cells of each of these kNNs. By repeatedly drawing random cells and computing an overlap distribution for each such kNN neighbourhood, we obtained a distribution of overlap widths which can then be compared to the overlap width of the smaller cluster of size *S*2. We also obtained similar results with alternative ways of dividing the clusters into smaller groups of cells, such as by sub-clustering the data into smaller clusters using standard clustering algorithms or by randomly subsampling cells (data not shown).

Secondly, as a negative control and to check that our results are not a generic feature due to common properties of RNA-seq data such as underlying count statistics, we simulated random uncorrelated cells. For this we used the *splatter* [50] package, which takes into account not only the negative binomial distribution of RNA-seq count data, but also additional features such as high expression outlier genes, differing sequencing depths (library sizes) between cells, and so on. We simulated 2000 cells and 10000 genes and verified from dimensionality reduction plots that there is no inherent structure in this data set (Fig. S3). We then computed fluctuations and overlaps in a similar manner and observed a near-perfect 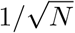 scaling for the overlap width with the number of genes (Fig. S2). We checked that these results remained consistent when we sub-clustered the data into smaller clusters, calculated the overlaps on kNN neighbourhoods or randomly subsampled cells (data not shown). Finally, we checked as previously that upon subsampling cells using kNN neighbourhoods, the overlap width does not depend noticeably on the number of subsampled cells (Fig. S2).

#### 1.2 Pan-GI Cell Atlas

Here we discussed the bioinformatics pipeline for Fig. 3.

To obtain a biologically meaningful and numerically well-conditioned gene set for model inference, we applied a unified gene-selection pipeline across all datasets. Raw counts were normalized to a constant library size and log-transformed, which mitigates cell-to-cell variability arising from sequencing depth and other global technical factors. While technical noise is inevitably present in single-cell measurements, it primarily affects the overall magnitude of fluctuations and is not expected to selectively induce population-specific correlation structures within the same dataset. We therefore focus on relative differences in covariance structure across populations, which are robust to such global sources of noise. We retained the top 10,000 highly variable genes to preserve broad transcriptional variation, including subtle regulatory programmes. For each annotated cell type, these HVGs were ranked using the Wilcoxon rank-sum statistic (implemented via the FindAllMarkers function in R), and the top 1,000 genes by this score were taken as a marker panel. This procedure enriches for genes that are both variable and discriminative for each cell type, while maintaining wide coverage of the underlying regulatory network. In this way, the resulting set of approximately 1,000 marker genes per cell type provides a broad and diverse sampling of transcriptional programmes under fixed parameter constraints, enabling the Boltzmann machine and covariance-based analyses to probe the underlying regulatory structure more effectively.

For model training on the Pan-GI Cell Atlas, we select the 100 top-ranked marker genes, balancing biological coverage with statistical tractability. Using 100 genes yields a parameter space of order 10^4^, comparable to the typical number of available datapoints (∼ 10^4^ cells), preventing overparameterization while retaining the dominant structure of gene-gene covariation. Expression values were centered and scaled, and the Boltzmann machine was fitted to this normalized data. The covariance matrix associated with the fitted model closely matched the empirical covariance, demonstrating that the gene-selection procedure preserves the essential structure of transcriptional fluctuations while enabling accurate inference of the interaction parameters. To assess whether the leading covariance eigenvalue *λ*_max_ probes the gene network more broadly than any specific marker choice, we additionally drew 100 random subsets of 100 genes from the 1000-gene marker panel for each Pan-GI cell type and computed *λ*_max_ on each subset, reporting the mean across these resamples. Because marker selection could in principle enrich for co-regulated genes, we verified that qualitative differences in *λ*_max_ persist across randomly sampled gene subsets, indicating that the observed coordination is not restricted to specific marker panels. For the zebrafish heart regeneration and human pulmonary disease analyses, we used the same Wilcoxon-based ranking and directly selected the top 100 marker genes per (cell type, time point) or (cell type, condition) as the basis for covariance and *λ*_max_ calculations.

In addition, for each cell type under investigation, defined in the main text as corresponding to a fixed point or steady state of the gene-expression dynamics, we selected cells that were closely located in gene-expression space to ensure that the modelled covariance reflects fluctuations around a coherent and well-defined dynamical state. For the Pan-GI atlas, we restricted our analysis to annotated cell types with at least 5000 cells and, where necessary, subsampled down to a maximum of 10000 cells per type by aggregating local neighbourhoods in the *k*-nearest-neighbour graph constructed during preprocessing. This strategy limits computational cost while preserving local coherence in expression space.

#### 1.3 Hearth regeneration and Pulmonary disease

Here we discussed the bionformatics pipeline for Fig. 4.

We applied the same feature-selection procedure described in Methods (Section 1.2), retaining the 100 most informative marker genes for each identified cluster. For the zebrafish heart regeneration scRNA-seq dataset, clusters were defined according to both cell type and time post-injury, consistent with the original study [44]. For the human pulmonary disease dataset, clusters were defined based on patient condition (disease versus control), following the annotation strategy of the original publication [45]. All p-values reported in Fig. 4b were computed using the Wilcox test. The p-values in Fig. 4d are computed with a bootstrapping method. For each cluster, we subsampled 50 times groups of 20 cells and computed the maximum eigenvalue of the corresponding covariance matrix. The distributions of maximum eigenvalues are shown in Fig. SS5b for each individual cluster with the numerical p-values of Fig. 4d.

##### Projection onto the leading covariance eigenmode

For each cluster, eigenvalue decomposition of the corresponding gene–gene covariance matrix *C* was performed according to

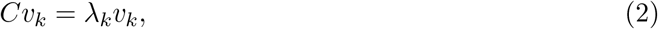

where *λ*_*k*_ and *v*_*k*_ denote the *k*-th eigenvalue and eigenvector, respectively, with eigenvalues ordered as *λ*_1_ ≥ *λ*_2_ ≥ ·· · ≥ *λ*_*p*_, where *p* denotes the number of genes included. The largest eigenvalue quantifies the strength of the dominant coordinated fluctuation mode within the cluster. To assess how individual cells align with this dominant mode, each cell’s centered expression vector 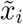 was projected onto the leading eigenvector *v*_1_ via the scalar projection

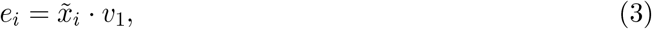

where *e*_*i*_ (referred to as the projected eigenvector) represents the amplitude of the dominant covariance mode for cell *i*, which we defined as eigenscore.

##### T-cell and macrophage score

The T-cell and macrophage score was build by first selecting typical markers in both cell types. T-cells markers: cd3d, cd247, lck, zap70, lat, trac, cd4, cd8a and Macrophages markers: mpeg1.1, csf1ra, marco, lyz, mrc1. We then use the build-in AddModuleScore() function in Seurat 4.4.1 on the selected markers.

## 2 Restricted Boltzmann machine analysis

In this section we provide details on the machine-learning algorithm and the bioinformatic pipeline supporting Fig. 3. We emphasize that the inferred parameters correspond to effective couplings, whose individual values are not directly interpreted biologically. Our aim is not to reconstruct specific gene–gene interactions, but to characterize statistical properties of the coupling matrix *W*_*ij*_, a proxy for the real couplings *J*_*ij*_, that reflect the collective organization of gene-expression fluctuations.

Because the number of inferred parameters scales quadratically with the number of genes (of order 10^4^ in our datasets), the model is necessarily high-dimensional relative to the available samples. Individual couplings are therefore not uniquely identifiable and may be subject to underdetermination. For this reason, we focus exclusively on robust aggregate quantities, such as the width of the coupling distribution, rather than on single interaction values.

### 2.1 Covariance matrix structure

The energy function of a Gaussian–Gaussian RBM is given by

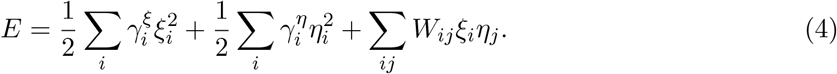

This energy function is formally equivalent to Eq. (1). The weights *W*_*ij*_ play the role of effective gene-gene interaction strengths *J*_*ij*_ in Eq. (S.5), up to diagonal terms. These diagonal contributions scale linearly with *N* and have a negligible effect on the inferred statistics compared to the off-diagonal elements, whose number scales as *N* ^2^. After fitting the RBM to the data, we therefore estimate the scaled standard deviation of gene–gene interactions, *σ*,

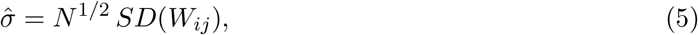

which provides a proxy for the true interaction variability *σ*. To compare the fitted model and data, we use the effective covariance matrix of the mRNA fluctuations obtained by integrating out the hidden states (namely the protein fluctuations). As the model is a Gaussian-Gaussian RBM, the integration over the ***ξ*** variables can be carried out exactly, yielding the following covariance matrix for the ***η***:

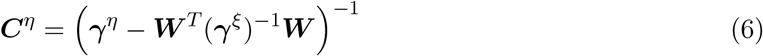

where *γ*^*ε*^ and *γ*^*ω*^ are diagonal matrices constructed from the corresponding coefficient vectors. This model covariance structure is used to compare model and data in Fig. 3 and Fig. S4.

### 2.2 RBM training set-up

Across all celltypes, we used a common strategy and same initial parameters to initialize the RBM training:

- The self-interaction coefficients of the protein fluctuations ***ξ*** are fixed to 1 and kept frozen, i.e. 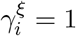. This is justified by the fact that any change in 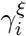 can be absorbed into a rescaling of the weights *W*_*ij*_.
- To ensure positivity, the self-interaction coefficients of the mRNA fluctuations ***η*** are reparametrized using a softplus function, i.e. 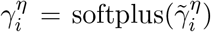 with softplus(*x*) = ln(1 + *e*^*x*^). The initial value for each gene *i* is set to 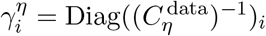.
- The interaction terms *W*_*ij*_ are initialized independently from a Gaussian distribution with zero mean and a standard deviation proportional to 1*/N*, where *N* is the number of genes. A single initialization scale was found to be insufficiently robust, so we performed a grid search on this hyper-parameter.
- We used mini-batches of size 200 for all datasets.
- The fitting procedure was carried out using persistent contrastive divergence with one Gibbs step [51].
- No regularization was applied during training. However, to avoid overfitting, we introduced early stopping criterion with a patience of 500 epochs, monitoring the difference between model and data energy losses, which serves as a Monte Carlo estimate of the negative log-likelihood of the observed dataset.

All code and data used for the RBM fitting are available at [39].

