## Supplementary Figures and Theory for "Coordinated gene expression variability encodes the regulatory state of cells": SI.pdf

### Supplementary Information

In this Supplementary information we derive the details of the theoretical methods in support of the main text. The startin point is the master equation describing the kinetics of mRNAs and protein in the gene regulatory networks which is given by,

$$\frac{\partial P}{\partial t} = \sum_{i=1}^{N} \left[ d_i \left(\mathcal{E}_i^1 - 1\right) n_i + \left(\mathcal{F}_i^1 - 1\right) \gamma_i m_i + \Omega \left(\mathcal{E}_i^{-1} - 1\right) g_i m_i + \Omega \left(\mathcal{F}_i^{-1} - 1\right) \sum_{j \in e(i)} f_{ji}^e \left(\frac{n_j}{\Omega}\right) \right] P \tag{S.1}$$

where $j \in e(i)$ signifies that the sum goes over all the transcription factors $j$ of gene $i$ and we used the notation of step operators $\mathcal{E}_i^{\pm} G\left(\mathbf{n}, \mathbf{m}\right) = G\left(\{\mathbf{n}, n_i \pm 1\}, \mathbf{m}\right)$, with $G$ an arbitrary function of protein and mRNA, and similarly for $\mathcal{F}_i^{\pm}$ which acts on the mRNA dependent part. In the following we are going to use eq. (S.1) in order to derive collective gene expression fluctuations dynamics.

#### A Distribution of the protein and mRNA fluctuations

In this section, we derive the "Hamiltonian" (1) starting from the master equation (S.1). To this end, we start by searching for the mean-field solutions describing average copy numbers of protein ($\phi$) and mRNA ($\psi$). This can be easily derive as in [1] and follows,

$$\begin{aligned} \frac{\partial \phi_i}{\partial t} &= g_i \psi_i - d_i \phi_i \,, \\ \frac{\partial \psi_i}{\partial t} &= -\gamma_i \psi_i + \sum_{j \in a(i)} f_{ji}^a \left(\phi_j\right) + \sum_{j \in r(i)} f_{ji}^r \left(\phi_j\right) \,, \end{aligned} \tag{S.2}$$

where $f_{ji}^r(\phi_j) = 1/(1 + \phi_j^{H_{ji}})$ and $f_{ji}^a(\phi_j) = \phi_j^{H_{ji}}/(1 + \phi_j^{H_{ji}})$ are the Hill functions for inihbitors and activators respectively where $H_{ji}$ is the Hill coefficient.

Systems of equation like (S.2) may exhibit multiple attractors which are defined by mRNA and protein concentrations such that $\partial_t \phi = \partial_t \psi_i = 0 \, \forall i = 1, \ldots, N$. In particular, close to one of those attractors we can expand Eq. (S.2) as $\phi_i = \phi_i^\star + \xi_i$ and $\psi_i = \psi_i^\star + \eta_i$ and we obtain,

$$\begin{aligned} \partial_t \xi_i &= g_i \eta_i - d_i \xi + \sqrt{D_i^n} W_i^\xi \,, \\ \partial_t \eta_i &= -\gamma_i \eta_i + \sum_{j \in e(i)} f'_{ji}(\phi_j^*) \xi_j + \sqrt{D_i^m} W_i^\eta \,, \end{aligned} \tag{S.3}$$

where $W^\xi$ and $W^\eta$ are unitary uncorrelated Gaussian white noises and $D_i^n$, $D_i^m$ are typical protein and mRNA fluctuations respectively. Here $\sum_{j \in e(i)}$ stands for the sum of the derivative of both activators and inhibitors. In the limit of fast degradation $\gamma_i \gg 1$ the two coupled equations reduce to a single equation for the protein fluctuations as the second equation becomes deterministic and can be integrated out.

$$\partial_t \xi_i = -d_i \xi_i + g_i \sum_{j \in e(i)} f'_{ji}(\phi_j^*) \xi_j / \gamma_j + \sqrt{D^n} W^\xi \,. \tag{S.4}$$

This fast degradation limit is valid in bacteria [2], but does not hold in eukaryotic systems. Here we look for approximate solutions of the distribution of protein and mRNA fluctuations as of the form, $\Pi = e^{-H}/Z$. We take the limit, $D_i^m \gg D_i^n$ which is expect as mRNA has more sources of fluctuations compared to protein, we get the approximate distribution,

$$\mathcal{H} = \sum_{i=1}^{N} \frac{d_i}{2D_i^n}\xi_i^2 + \sum_{i=1}^{N} \frac{\gamma_i}{2D_i^m}\eta_i^2 - \sum_{i=1}^{N} \frac{2g_i}{D_i^n}\xi_i\eta_i + \sum_{i,j=1}^{N} J_{ij}\xi_j\eta_i \tag{S.5}$$

where $J_{ij} = f'_{ji}(\phi_j^*)/D_i^m$. Eq. (S.5) is approximate as the term in $\xi_i\eta_i$ would give rise to a contribution in the Langevin dynamics of $\xi_i$ and $\eta_j$ which are not originally present in (S.3), namely one would get

$$\begin{aligned} \partial_t\xi_i &= g_i\eta_i - d_i\xi_i + \sum_{j\in e(i)} \frac{f'_{ji}(\phi_j^*)D_i^n}{D_i^m}\eta_j + \sqrt{D^n}W^\xi \\ \partial_t\eta_i &= -\gamma_i\eta_i + \sum_{j\in e(i)} f'_{ji}(\phi_j^*)\xi_j + g_i\frac{D_i^m}{D_i^n}\xi_i + \sqrt{D^m}W^\eta\,. \end{aligned} \tag{S.6}$$

As fluctuations in mRNA are dominant then $D_i^m \gg D_i^n$, then the first equation is reduced to the exact Langevin equation (S.3), while the second one has still a term in $\xi_i$ which is not originally present. We can however safely reabsorb the latter by rescaling the term in $\sum_{j\in e(i)} f'(\phi_j^*)\xi_j$ (corresponding to self-activation or repression) in such a way that the dynamics is fully described by the Hamiltonian (S.5).

### B Discussion on model assumptions

In the following, we consider that the main source of heterogeneity arises from the derivative of the Hill-functions such that we take the couplings $J_{ij}$ to be Gaussian distributed with vanishing mean and standard deviation $\sigma/\sqrt{N}$. This choice is in accordance with previous studies [3]. Despite gene regulatory networks are sparse, meaning that there are only few non-zero couplings, in [4] it is shown for a very similar model that dilution, even when extreme, does not change qualitatively the phase diagram. From now on, we will leverage the restriction on the sparseness of the couplings for the sake of theoretical tractability. All the other parameters are taken to be constant and independent on the gene $i$. With this parameter choice, the "Hamiltonian" in Eq. (1) is identical to the one of a bipartite, soft spin-glass with asymmetric couplings of the form $J_{ij}$ [5, 6]. Here, we reason that fluctuations can not be Gaussian unconstrained for mainly two reasons. First, unconstrained fluctuations for every gene products are biologically unreasonable and second, unconstrained fluctuations could bring the system toward another steady state. As we are interested only in fluctuations around a given stationary state, in the following we will consider theoretically two main classes of constraints: a spherical constraints, $\sum_i \xi_i^2 = N$ and a binary ine $\xi_i = \pm 1$ and similarly for $\eta_i$. Binary constraints have been widely used in boolean approximations to gene networks [7]. Spherically constrained fluctuations have the advantage to be theoretically simpler to study and in some limits to give results comparable to the binary case. In the main text we found that our Hamiltonian Eq. (S.5) is identical to the one of a RBM. Here we note that RBMs have been found to be under certain conditions, the limit of Hopfield neural networks [8, 9], which have been recently used to infer attractors of developmental landscapes [10, 11, 12]. Here we note, that while in these studies the focus was to few attractors, such that the Hopfield model is the very well behaved as it encodes different limited patterns; here we are interested in the collective fluctuations, such that the assumptions to have only few patterns does not hold. Indeed, in the limit of weakly correlation between the number of possible stored patterns $P$ and for numbers $P \sim \alpha N$, the Hopfield model maps exactly to the RBMs and so to our theory [8].

### C Derivation of the bipartite spin glass

In this section we show how we can use methods from spin-glass theory to study fluctuations of GRNs. The starting point is Eq. (1) of the main text,

$$\mathcal{H} = \sum_{i=1}^{N} V^{\xi}(\xi_i) + \sum_{i=1}^{N} V^{\eta}(\eta_i) + \sum_{ij} J_{ij}\xi_i\eta_j + \sum_{i=1}^{N} K_i^{\xi\eta}\xi_i\eta_i\,, \tag{S.7}$$

with $J_{ij}$ distributed according to a Gaussian with zero mean and standard deviation $\theta$, $N(0,\theta)$. To ensure the extensivity of the Hamiltonian in the limit $N \to \infty$ we define the standard deviation as $\theta = \sigma\big/\sqrt{N}$. Further, we set $V^{\xi}(\xi_i) = K^{\xi_i}\ \xi_i^2/2$ and similarly for $V^{\eta}$. To obtain results that are independent of the specific choice of couplings, we perform the average over the quenched average

$$[Z^n] = \int D\left[J_{ij}\right] D\left[\xi\right] D\left[\eta\right] P(J_{ij}) \exp(-\mathcal{H}_r)\,, \tag{S.8}$$

where $\mathcal{H}_r$ is the replicated Hamiltonian. The off-diagonal terms in the partition function,

$$\int \prod_{ij} \mathrm{d}J_{ij} e^{-\frac{J_{ij}^2}{2W^2}} e^{-\sum_a J_{ij}\xi_i^a\eta_j^a}\,, \tag{S.9}$$

have to be evaluated with a Gaussian integration over the couplings $J_{ij}$. After doing so the quenched average partition functions is,

$$\begin{aligned}[Z^n] = \mathrm{Tr}_n \exp\Big[\frac{\theta^2}{2} \sum_{a,b,i,j} \xi_i^a\xi_i^b\eta_j^a\eta_j^b + \sum_{i,a} V^{\xi}(\xi_i^a) \\ + \sum_{j,a} V^{\eta}(\eta_j^a) + \sum_{i,a} K^{\xi\eta}\xi_i^a\eta_i^a\Big]\,,\end{aligned} \tag{S.10}$$

where $\mathrm{Tr}_n$ in case of non-binary variables is and integrale over the variables and it is the trace when $\xi_i, \eta_i = \pm 1$.

To simplify the quartic term in Eq.(S.10) we further introduce the integral transformation

$$e^{\frac{BC}{\sqrt{2}\omega}} \sim \int e^{-\omega(x^2-\sqrt{2}xy+y^2+\frac{1}{2}\tilde{x}^2+\frac{1}{2}\tilde{y}^2)} e^{B(x+i\tilde{x})+C(y+i\tilde{y})}. \tag{S.11}$$

Inserting this transformation to the Hamiltonian with the following variables

$$\omega = N\frac{\sigma^2}{2\sqrt{2}}\,, \qquad B = \frac{\sigma^2}{2}\sum_i \xi_i^a\xi_i^b\,, \qquad C = \frac{\sigma^2}{2}\sum_j \eta_j^a\eta_j^b\,, \tag{S.12}$$

the quenched average partition function is

$$
\begin{aligned}
[Z^n] =& \int \prod_{a,b} \mathrm{d}x_{ab} \mathrm{d}\tilde{x}_{ab} \mathrm{d}y_{ab} \mathrm{d}\tilde{y}_{ab} e^{-NnF_n} , \\
nF_n =& \frac{\sigma^2}{2} \sum_{a \neq b} \left( \frac{x_{ab}^2}{\sqrt{2}} + \frac{y_{ab}^2}{\sqrt{2}} + \frac{\tilde{x}_{ab}^2}{2\sqrt{2}} + \frac{\tilde{y}_{ab}^2}{2\sqrt{2}} - x_{ab} y_{ab} \right) \\
&+ \frac{\sigma^2}{2} \sum_{a} \left( \frac{x_{aa}^2}{\sqrt{2}} + \frac{y_{aa}^2}{\sqrt{2}} + \frac{\tilde{x}_{aa}^2}{2\sqrt{2}} + \frac{\tilde{y}_{aa}^2}{2\sqrt{2}} - x_{aa} y_{aa} \right) \\
&- \log \mathrm{Tr}_{n\xi\eta} \Psi_\xi \Psi_\eta \Psi_{\eta\xi} ,
\end{aligned}
\tag{S.13}
$$

where

$$
\begin{aligned}
\Psi_\xi &= \exp \left[ \sum_{a,b} \frac{\sigma^2}{2} \left( x_{ab} + i\tilde{x}_{ab} \right) \xi^a \xi^b + \sum_{a,i} V^\xi(\xi^a) + \right] \\
\Psi_\eta &= \exp \left[ \sum_{a,b} \frac{\sigma^2}{2} \left( y_{ab} + i\tilde{y}_{ab} \right) \eta^a \eta^b + \sum_{a,i} V^\eta(\eta^a) \right] \\
\Psi_{\eta\xi} &= \exp \left[ \sum_{a} K^{\xi\eta} \xi^a \eta^a \right] .
\end{aligned}
\tag{S.14}
$$

As the the exponential term carries a factor $N$ we can evaluate the trace and the saddle point equations in the limit $N \to \infty$. To do so, we perform a change of variables (we already summed over $i$ neglecting variability in local parameters)

$$
\begin{aligned}
Q_{ab}^\xi = \left( x_{ab} + i\tilde{x}_{ab} \right) \; Q_{ab}^\eta = \left( y_{ab} + i\tilde{y}_{ab} \right) , \\
\hat{Q}_{ab}^\xi = \left( x_{ab} - i\tilde{x}_{ab} \right) \; \hat{Q}_{ab}^\eta = \left( y_{ab} - i\tilde{y}_{ab} \right) .
\end{aligned}
\tag{S.15}
$$

Here we explicit the diagonal components $Q_{aa}$ for reasons that will become clear in the next section. Rewriting the free energy and performing the saddle point over the hatted variables (which do not enter into the traces) the resulting free energy is

$$
nF_n = \frac{\sigma^2}{2} \sum_{a \neq b} Q_{ab}^\xi Q_{ab}^\eta + \frac{\sigma^2}{2} \sum_{a} Q_{aa}^\xi Q_{aa}^\eta - \log \mathrm{Tr}_{n\eta\xi} \Psi \tag{S.16}
$$

where $\Psi = \Psi_\eta \Psi_\xi \Psi_{\eta\xi}$.

### D Replica symmetric solution

In this section we provide detailed calculation leading to the phase diagram Fig. 1e. Starting from Eq. (S.16), we consider the replica symmetric ansatz

$$
\begin{aligned}
Q_{ab}^\xi &= q_0^\xi , \qquad Q_{aa}^\xi = q_D^\xi , \\
Q_{ab}^\eta &= q_0^\eta , \qquad Q_{aa}^\eta = q_D^\eta .
\end{aligned}
\tag{S.17}
$$

Substituting this ansatz into the free energy leads to

$$nF_n = \frac{\sigma^2}{2}\sum_{a\neq b}\left(q_0^\xi q_0^\eta\right) + \frac{\sigma^2}{2}\sum_a \left(q_D^\xi q_D^\eta\right) - \log \mathrm{Tr}_{n\eta\xi}\Psi\,, \tag{S.18}$$

The replicated quenched averaged partition function is

$$\begin{aligned}[Z^n] = \int dq_0^\xi dq_0^\eta dq_D^\xi dq_D^\eta \exp\left[N\left(\sigma^2 q_0^\xi q_0^\eta \frac{n(n-1)}{2}\right)\right] \\ \exp\left[\frac{\sigma^2}{2} n q_D^\xi q_D^\eta - \left(\log\int\prod_a d\xi^a d\eta^a \Psi\right)\right]\end{aligned} \tag{S.19}$$

where we made $Tr$ is made explicit for non-binary variable. In the previous equation we observe that both $\Psi_\xi$ and $\Psi_\eta$ factorize , such that the only cumbersome term to evaluate is,

$$\begin{aligned}\int\prod_a d\xi^a \exp\left[\frac{\sigma^2}{2}q_0^\xi\left(\sum_a \xi^a\right)^2\right] \\ \exp\left[\frac{\sigma^2}{2}\left(q_D^\xi - q_0^\xi\right)\sum_a(\xi^a)^2 + \sum_a V^\xi(\xi^a)\right]\,.\end{aligned} \tag{S.20}$$

The previous integral can be easiliy calculated upon performing one more Hubbard Stratonovich transformation for the variable $S = \sum_a \xi_a$. By doing so the integral to calculate reduces to

$$\int dz e^{-z^2/2}\int\prod_a d\xi^a \exp\left[-\sum_a H_{RS}^\xi(\xi_a, z)\right]\,, \tag{S.21}$$

with

$$\mathcal{H}_{RS}^\xi(\xi^a) = -\sigma\sqrt{q_0^\xi}z\xi^a - \frac{\sigma^2}{2}\left(q_D^\xi - q_0^\xi\right)(\xi^a)^2 + V^a(\xi^a)\,, \tag{S.22}$$

and similarly for $\mathcal{H}_{RS}^\eta$.
Taken together the free energy to minimize is,

$$nF_n = \frac{\sigma^2}{2}n(n-1)q_0^\xi q_0^\eta + \frac{\sigma^2}{2}nq_D^\xi q_D^\eta - \log \mathrm{Tr}\Psi\,. \tag{S.23}$$

The previous equations might be hard to evaluate for generic spin-systems. Here, we make the assumptions that, as fluctuations are typically of order $\sqrt{N}$, we can add a spherical constraint to the previous equation

$$\frac{1}{N}\sum_i \xi_i^2 = 1\,, \qquad\qquad \frac{1}{N}\sum_i \eta_i^2 = 1\,, \tag{S.24}$$

effectively restricting the fluctuations to an hypersphere. This spherical constraints implies $q_D = 1$. We then add $2nN$ Lagrange multipliers $(\lambda_a^\xi, \lambda_a^\eta)$ in (S.19) using the integral representation of the delta functions

$$1 = \int d\xi_i^a \delta\left(\sum_i(\xi_i^a)^2 - N\right)\,, \tag{S.25}$$

and similarly for $\eta$. The constrained free energy hence simplifies to,

$$
\begin{aligned}
nF_n =& \frac{\sigma^2}{2} n(n-1) q_0^\xi q_0^\eta \\
&+ \frac{\sigma^2}{2} n + \sum_a (\lambda_a^\xi + \lambda_a^\eta) - (\log \mathrm{Tr}\Psi) \,,
\end{aligned}
\tag{S.26}
$$

where

$$
\mathcal{H}_{RS}^\xi = -\sigma\sqrt{q_0^\xi} z\xi^a - \frac{\sigma^2}{2}\left(1-q_0^\xi\right)(\xi^a)^2 + V^a(\xi^a) - \lambda_a^\xi (\xi^a)^2 \tag{S.27}
$$

and similarly for $\mathcal{H}_{RS}^\eta$. Moreover, $V(\xi_a)$ and $V(\eta_a)$ are quadratic in $\eta_a, \xi_a$ and their sum over replicas is a constant that we can safely neglect for the future computations. Consistently, we consider the replica symmetric ansatz for the multipliers: $\lambda_a^\xi = \lambda_0^\xi$ , $\lambda_a^\xi = \lambda_0^\eta$ .

#### D I Overlaps of spherically constrained fluctuations

In order to find the value of the overlaps that minimize the free energy we take the saddle point equations

$$
\frac{\delta F}{\delta q_0^\xi} = 0 \,, \qquad \frac{\delta F}{\delta \lambda_0^\xi} = 0 \,, \qquad \frac{\delta F}{\delta q_0^\eta} = 0 \,, \qquad \frac{\delta F}{\delta \lambda_0^\eta} = 0 \,. \tag{S.28}
$$

These equations reduce to the integral solution of four coupled equations, namely

$$
\begin{aligned}
q_0^\xi &= \frac{\int DzDw \prod^c d\xi^c d\eta^c \xi^a \xi^b e^{-\sum_c \mathcal{H}_{RS}(\xi^c,\eta^c,z)}}{\int DzDw \prod_c d\xi^c d\eta^c e^{-\sum_c H_{RS}(\xi^c,\eta^c,z)}} \,, \\
1 &= \frac{\int DzDw \prod_c d\xi^c d\eta^c (\xi^a)^2 e^{-\sum_c \mathcal{H}_{RS}(\xi^c,\eta^c,z)}}{\int DzDw \prod_c d\xi^c d\eta^c e^{-\sum_c H_{RS}(\xi^c,\eta^c,z)}} \,, \\
q_0^\eta &= \frac{\int DzDw \prod_c d\xi^c d\eta^c \eta^a \eta^b e^{-\sum_c \mathcal{H}_{RS}(\xi^c,\eta^c,z)}}{\int DzDw \prod_c d\xi^c d\eta^c e^{-\sum_c H_{RS}(\xi^c,\eta^c,z)}} \,, \\
1 &= \frac{\int DzDw \prod_c d\xi^c d\eta^c (\eta^a)^2 e^{-\sum_c \mathcal{H}_{RS}(\xi^c,\eta^c,z)}}{\int DzDw \prod_c d\xi^c d\eta^c e^{-\sum_c H_{RS}(\xi^c,\eta^c,z)}} \,,
\end{aligned}
\tag{S.29}
$$

where we defined $Dz = dze^{-\frac{z^2}{2}}$, $Dw = dwe^{-\frac{w^2}{2}}$ and

$$
\begin{aligned}
H_{RS} =& - z\sigma\sqrt{q_0^\xi}\xi^a - w\sigma\sqrt{q_0^\eta}\eta^a + K^{\xi\eta}\eta^a\xi^a \\
&- \sigma^2\left(1-q_0^\xi\right)\frac{(\xi^a)^2}{2} - \sigma^2\left(1-q_0^\eta\right)\frac{(\eta^a)^2}{2} \\
&+ \lambda_0^\xi(\xi^a)^2 + \lambda_0^\eta(\eta^a)^2 \,.
\end{aligned}
\tag{S.30}
$$

In the limit of strong interactions ($\sigma \to \infty$) we can solve the internal integral with the saddle point method. We then replace the integral with $\xi_a^*, \eta_a^*$ that satisfies the equations $\frac{\partial \mathcal{H}_{RS}}{\partial \xi_a}|_{\xi_a^*,\eta_a^*} = 0$, $\frac{\partial \mathcal{H}_{RS}}{\partial \eta_a}|_{\xi_a^*,\eta_a^*} = 0$. In case of RS solution we simplify to

$$
q_0^\xi = \int DzDw\, \eta_{a^2}^* \,. \tag{S.31}
$$

As an example, when $K^{\xi\eta}=0$

$$q_0^\xi = \left(\frac{\sigma\sqrt{q_0}}{\sigma^2(1-q_0)}\right)^2 \int dz e^{-\frac{z^2}{2}} z^2\,, \tag{S.32}$$

with the known solution [13]

$$q_0^\xi = 1-\frac{1}{\sigma} = q_0^\eta\,. \tag{S.33}$$

The replica symmetric Hamiltonian has the symmetry under the exchange $\xi \longleftrightarrow \eta$, which simplifies the next computation as it implies that $q_0^\xi = q_0^\eta = q_0$. The exact solution of the coupled equations for the overlap and Lagrange multiplier can be recasted as [14],

$$\begin{aligned} q_0^\xi &= \int DzDw \left(\frac{\int d\xi^a d\eta^a \xi^a e^{-\sum_c \mathcal{H}_{RS}(\xi^c,\eta^c,z)}}{\int d\xi^a d\eta^a e^{-H_{RS}(\xi^a,\eta^a,z)}}\right)^2 \\ 1 &= \int DzDw \frac{\int d\xi^a d\eta^a {\xi^a}^2 e^{-\sum_c \mathcal{H}_{RS}(\xi^c,\eta^c,z)}}{\int d\xi^a d\eta^a e^{-H_{RS}(\xi^a,\eta^a,z)}}\,, \end{aligned} \tag{S.34}$$

and the solution is

$$q_0 = 1 - \frac{\frac{1}{\sigma}\left(\sqrt{\frac{8}{\sigma^2}(K^{\xi\eta})^2+1}+3\right)}{2\sqrt{2}\sqrt{\frac{2}{\sigma^2}(K^{\xi\eta})^2+\sqrt{\frac{8}{\sigma^2}(K^{\xi\eta})^2+1}+1}}\,. \tag{S.35}$$

Moreover, we found consistently that $\overline{\langle\xi_i\rangle}=0$, where $\overline{(\cdot)}$ indicates the average over the disorder $J_{ij}$. Indeed,

$$\overline{\langle\xi_i\rangle} = \frac{\int DzDw \prod_c d\xi_c d\eta_c \xi_a e^{-\sum_c \mathcal{H}_{RS}(\xi_c,\eta_c,z)}}{\int DzDw \prod_c d\xi_c d\eta_c e^{-\sum_c \mathcal{H}_{RS}(\xi_c,\eta_c,z)}} \tag{S.36}$$

in the limit $\sigma\to 0$, we substitute the internal integral with the saddle point and giving that $\xi^*$ is an odd function of $z, w$ the result of the external integral is zero. The same results apply to $\overline{\langle\eta_i\rangle}$. This is clearly in agreement with our choice of working with fluctuations only. In the next section we are going to show an alternative derivation.

#### D II Alternative calculation

An alternative way to calculate the overlaps could be to start from (S.13) and, following [15], to insert the definition of the overlaps and the spherical constraints as delta distributions with Lagrange multipliers $\lambda_{ab}$. The resulting free energy is

$$\begin{aligned} nF_n =& \frac{\sigma^2}{2}\sum_{a,b} q_{ab}^\xi q_{ab}^\eta + \sum_{ab}\lambda_{ab}^\xi q_{ab} + \sum_{ab}\lambda_{ab}^\eta q_{ab} + \\ & - \log \mathrm{Tr}_{n\eta\xi}\Psi, \end{aligned} \tag{S.37}$$

where

$$\begin{aligned} \mathrm{Tr}_{n\eta\xi}\Psi = \int d\boldsymbol{\xi} d\boldsymbol{\eta} \exp&\left\{\left[\sum_{a,b}\lambda_{ab}^\xi \xi^a\xi^b\right]\right\} \\ \exp&\left\{\left[\sum_{a,b}\lambda_{ab}^\eta \eta^a\eta^b + \sum_a K^{\xi\eta}\right]\right\}. \end{aligned} \tag{S.38}$$

We omitted the potential due to the spherical constraint. The argument of the exponential in the last expression can be rewritten as

$$(\boldsymbol{\xi}, \boldsymbol{\eta}) \begin{pmatrix} \boldsymbol{\lambda}^{\boldsymbol{\eta}} & \frac{K^{\xi\eta}}{2}\boldsymbol{I} \\ \frac{K^{\xi\eta}}{2}\boldsymbol{I} & \boldsymbol{\lambda}^{\boldsymbol{\eta}} \end{pmatrix} \begin{pmatrix} \boldsymbol{\xi} \\ \boldsymbol{\eta} \end{pmatrix} , \tag{S.39}$$

and the free energy becomes

$$\begin{aligned} nF_n =& \frac{\sigma^2}{2} \sum_{a,b} q^{\xi}_{ab} q^{\eta}_{ab} + \sum_{ab} \lambda^{\xi}_{ab} q_{ab} + \sum_{ab} \lambda^{\eta}_{ab} q_{ab} \\ & - \frac{1}{2} \log\det\left( \boldsymbol{\lambda}^{\boldsymbol{\eta}} \boldsymbol{\lambda}^{\boldsymbol{\xi}} - \frac{\tilde{K^2}_{\xi,\eta}}{4} \boldsymbol{I} \right) . \end{aligned} \tag{S.40}$$

When $K^{\xi\eta} \gg 1$ (ignoring constants) the last term of the previous equation is approximated as,

$$\log\det\left[ \boldsymbol{\lambda}^{\boldsymbol{\eta}} \boldsymbol{\lambda}^{\boldsymbol{\xi}} - \frac{(K^{\xi\eta})^2}{4} \boldsymbol{I} \right] \approx \mathrm{Tr}(\boldsymbol{\lambda}^{\boldsymbol{\eta}} \boldsymbol{\lambda}^{\boldsymbol{\xi}} \frac{4}{(K^{\xi\eta})^2}) . \tag{S.41}$$

In order to find the value of the overlaps that minimize the free energy we take the saddle point equations

$$\frac{\delta F}{\delta q^{\xi}_{ab}} = 0 , \quad \frac{\delta F}{\delta \lambda^{\xi}_{ab}} = 0 , \quad \frac{\delta F}{\delta q^{\eta}_{ab}} = 0 , \quad \frac{\delta F}{\delta \lambda^{\eta}_{ab}} = 0 . \tag{S.42}$$

The extremization with respect to $\lambda$ leads to

$$q^{\xi}_{ab} = \frac{2}{(K^{\xi\eta})^2} \lambda^{\eta}_{ba} , \ \xi \leftrightarrow \eta , \tag{S.43}$$

and

$$\frac{\sigma^2}{2} q^{\xi}_{ab} - q^{\eta}_{ab} = 0 , \ \xi \leftrightarrow \eta . \tag{S.44}$$

A part from the paramagnetic solution $q_{ab} = 0, a, b = 1 \dots n$, the last equation is always solved if $\sigma = \sqrt{2}$. The calculation can be carried similarly for $K^{\xi\eta} \ll 1$ and we found a limiting value $\sigma = 1$, as in the previous derivation.

#### D III Overlaps of binary fluctuations

In case of binary fluctuations the free energy reduces to the one of a bipartite Sherrington-Kirkpatrick model -where the spherical constraint is automatically satisfied- with an additional term that couples $\xi_i$ and $\eta_i$, namely $K^{\xi\eta}$,

$$nF_n = \frac{\sigma^2}{2} \sum_{a \neq b} \left( Q^{\xi}_{ab} Q^{\eta}_{ab} \right) + \frac{\sigma^2}{2} n - \log \mathrm{Tr}_{n\eta\xi} \Psi , \tag{S.45}$$

where again $\Psi = \Psi_\xi \Psi_\eta \Psi_{\eta,\xi}$ and

$$\begin{aligned} \Psi_\xi &= \exp\left[ \sum_{a \neq b} \frac{\sigma^2}{2} Q^{\xi}_{ab} \xi_a \xi_b \right] , \\ \Psi_\eta &= \exp\left[ \sum_{a \neq b} \frac{\sigma^2}{2} Q^{\eta}_{ab} \eta_a \eta_b \right] , \\ \Psi_{\eta\xi} &= \exp\left[ \sum_{a} K^{\xi\eta} \xi_a \eta_a \right] . \end{aligned} \tag{S.46}$$

We perform again a Hubbard-Stratonovich transformation and taking the RS solution we arrive to ($f = F_n$),

$$f = -\frac{\sigma^2}{2}(q_0^\xi - 1)(q_0^\eta - 1) - \langle [4\cosh\left(\sigma\sqrt{q_0^\xi}z\right)\cosh(\sigma\sqrt{q_0^\eta}w)\cosh\left(K^{\xi\eta}\right) + 4\sinh\left(\sigma\sqrt{q_0^\xi}z\right)\sinh\left(\sigma\sqrt{q_0^\eta}w\right)\sinh(K^{\xi\eta})]\rangle_{zw}\,. \tag{S.47}$$

Upon finding the saddle point as done previously and expanding the self-consistency equations, we find that the phase boundary between the paramagnetic and glassy phase is given by

$$1 + \tanh\left(K^{\xi\eta}\right) = 1/\sigma^2\,. \tag{S.48}$$

### E Path integral of spin glass dynamics

In this section we derive a simpler form for the dynamics of protein and mRNA fluctuations which will be useful for the calculations of dynamical auto-correlation functions. To this end, following [16, 17] we introduce a generating functional for the coupled Langevin equations (S.3),

$$Z[\boldsymbol{h}^\xi, \boldsymbol{h}^\eta] = \int \mathrm{D}[\boldsymbol{\xi}]\mathrm{D}[\boldsymbol{\eta}]\mathrm{D}[\hat{\boldsymbol{\xi}}]\mathrm{D}[\hat{\boldsymbol{\eta}}]e^{i\int dt \sum_j \left(h_j^\xi \xi_j + h_j^\eta \eta_j\right)} e^{i\int dt \sum_j \hat{\xi}_j\left[\partial_t \xi_j - g_j\eta_j + d_j\xi - D_j^n \hat{\xi}_j\right]} e^{i\int dt \sum_j \hat{\eta}_j\left[\partial_t \eta_j + \gamma_j\eta_j - \sum_{k\in e(j)} f'_{k,j}(\phi_k^*)\xi_k - \gamma_j b_j - D_j^m \hat{\eta}_j\right]}\,. \tag{S.49}$$

We initially isolate the part which includes disorder by replacing $G_j = \sum_{k\in e(j)} f'(\phi_k^*)\xi_k$. By doing so, we formally introduce a new delta

$$Z[\boldsymbol{h}^\xi, \boldsymbol{h}^\eta] = \int \mathrm{D}[\boldsymbol{\xi}]\mathrm{D}[\boldsymbol{\eta}]\mathrm{D}[\hat{\boldsymbol{\xi}}]\mathrm{D}[\hat{\boldsymbol{\eta}}]\mathrm{D}[\hat{\boldsymbol{G}}]\mathrm{D}[\boldsymbol{G}] e^{i\int dt \sum_j \left(h_j^\xi \xi_j + h_j^\eta \eta_j\right)} e^{i\int dt \sum_j \hat{G}_j^\xi\left(G_j - \sum_{k\in e(j)} f'_{k,j}(\phi_k^*)\xi_k\right)} e^{i\int dt \sum_j \hat{\xi}_j\left[\partial_t \xi_j - g_j\eta_j + d_j\xi - D_j^n \hat{\xi}_j\right]} e^{i\int dt \sum_j \hat{\eta}_j\left[\partial_t \eta_j + \gamma_j\eta_j - G_j - \gamma_j b_j - D_j^m \hat{\eta}_j\right]}\,. \tag{S.50}$$

In the dynamical representation of glassy fluctuations time plays a similar role as the replica index in the equilibrium regime. We then integrate over the couplings $f'_{k,j}(\phi_k^*) = J_{kj}$ with statistics, $\overline{J_{kj}} = 0$, $\overline{J_{kj}^2} = \sigma^2/N$, and $\overline{J_{kj}J_{jk}} = \lambda\sigma^2/N$. We further integrate over $J_{kj}$ ($\overline{\cdots}$) in the exponential

$$\overline{e^{-i\sum_j \int dt \hat{G}_j(t)\sum_{k\in e(j)} J_{kj}\xi_k(t)}}\,, \tag{S.51}$$

which results in

$$
\begin{aligned}
& e^{-\frac{\sigma^2}{2}N\int dtdt'\left(L(t,t')C^{\xi}(t,t')+\lambda K(t,t')K(t',t)\right)}\,, \\
& L(t,t') = \frac{1}{N}\sum_j \hat{G}_j(t)\hat{G}_j(t')\,, \\
& C^{\xi}(t,t') = \frac{1}{N}\sum_j \xi_j(t)\xi_j(t')\,, \\
& K(t,t') = \frac{1}{N}\sum_j \xi_j(t)\hat{G}_j(t')\,.
\end{aligned} \tag{S.52}
$$

From now on we can follow [18] and obtain the following simplified (in terms of number of equations) dynamics,

$$
\begin{aligned}
\partial_t \xi_i =& g_i\eta_i - d_i\xi_i + \sqrt{D_i^n}W_i^{\xi}\,, \\
\partial_t \eta_i =& -\gamma_i\eta_i + \sigma^2\lambda\int \mathrm{d}t'\chi_{\eta_i}^{\xi_i}(t,t')\xi_i(t') \\
& + \sigma W_c^{\eta_i} + \sqrt{D_i^m}W_i^{\eta}\,.
\end{aligned} \tag{S.53}
$$

$W_i^{\eta}$ and $W_i^{\xi}$ are Gaussian uncorrelated white noises with zero mean and unit variance and $\langle W_c^{\eta_i}(t)W_c^{\eta_i}(t')\rangle = C_i^{\xi}(t,t')$, $\overline{J_{ij}} = 0$, $\overline{J_{ij}^2} = \sigma^2/N$, and $\overline{J_{ij}J_{ji}} = \lambda\sigma^2/N$. $C_i^{\xi}(t,t')$ and $\chi_{\eta_i}^{\xi_i}(t,t')$ are respectively the auto-correlation and response functions of the protein fluctuations. As the equations factorize in the number of genes, we can compute individual gene autocorrelation and response functions.

### F Out of equilibrium dynamics of p-spin spherical asymmetric bipartite spin glasses

In this section we derive the behaviour of correlation functions of protein and mRNAs fluctuations. To do so, we write the dynamics of protein and mRNA fluctuations with a spherical constraint $\sum_i \xi_i(t)^2 = N\,, \sum_i \eta_i(t)^2 = N$ [19, 20, 21] upon introducing Lagrange multipliers $\mu^{\xi}(t), \mu^{\eta}(t)$,

$$
\begin{aligned}
\partial_t \xi_i(t) &= g_i\eta_i(t) - d_i\xi(t) - \mu_i^{\xi}(t)\xi(t) + \sqrt{D_i^n}W_i^{\xi}(t) \\
\partial_t \eta_i(t) &= -\gamma_i\eta_i(t) - \mu_i^{\eta}(t)\eta(t) + W_c^{\eta_i}(t)\,,
\end{aligned} \tag{S.54}
$$

with $\langle W_c^{\eta_i}(t)W_c^{\eta_i}(t')\rangle = D_i^m + \sigma^2 C_i^{\xi}(t,t')$.

From Eq. (S.54) we obtain the equations for the dynamics of autocorrelation functions,

$$
\begin{aligned}
\partial_t C^{\xi_i} &= g_i C^{\xi\eta} - d_i C_i^{\xi} - \mu^{\xi}(t)C_i^{\xi} + D_i^n\langle W_i^{\xi}(t)\xi_i(t')\rangle \\
\partial_t C_i^{\eta} &= -\gamma_i C_i^{\eta} - \mu^{\eta}(t)C_i^{\eta} + \langle W_c^{\eta_i}(t)\eta_i(t')\rangle\,,
\end{aligned} \tag{S.55}
$$

with $C_i^{\xi\eta} = \langle \xi_i(t)\eta_i(t')\rangle$ and for sake of notation we remove the dependencies on $t, t'$ of $C^{\xi\eta}$.

Starting from Eq. (S.55),

$$
\begin{aligned}
\partial_t C^{\xi} &= g_i C_i^{\xi\eta} - (d_i + \mu^{\xi})C_i^{\xi} + D_i^n\langle W^{\xi}(t)\xi(t')\rangle\,, \\
\partial_t C_i^{\eta} &= -\gamma_i C_i^{\eta} - \mu^{\eta_i}(t)C_i^{\eta} + \langle W_c^{\eta_i}(t)\eta(t')\rangle\,,
\end{aligned} \tag{S.56}
$$

the spherical constraint imposes $C_i^{\xi/\eta}(t,t')|_{t'\to t} = 1$. $\mu_i^{\xi/\eta}(t)$ can be evaluated by the relationship $\partial_t C_i^{\xi/\eta}(t,t')|_{t'\to t} + \partial_{t'} C_i^{\xi/\eta}(t,t')|_{t\to t'} = 1$

$$\begin{aligned}\mu^\xi(t) &= (g_i C_i^{\xi\eta}(t,t) - d_i) + D_i^n \langle W_i^\xi(t)\xi_i(t)\rangle\,,\\ \mu^\eta(t) &= -\gamma_i + \langle W_c^{\eta_i}(t)\eta_i(t)\rangle\,.\end{aligned} \tag{S.57}$$

We are then just left to compute the terms in the brackets. In particular with Novikov's theorem [22] the first one is

$$\langle W_i^\xi(t)\xi_i(t')\rangle = \chi_i^\xi(t,t')\,, \tag{S.58}$$

where the propagator $\chi_i^\xi(t,t') = \langle \frac{\delta\xi_i(t)}{\delta W_i^\xi(t')}\rangle$. We take $t' > t$ and so $\chi(t,t') = 0$ for $t' > t$ due to causality and $\lim_{t'\to t}\chi(t,t') = 1$. We then find $\mu_i^\xi(t) = g_i C_i^{\xi\eta}(t,t) - d_i + \frac{D_i^n}{2}$. The second average is

$$\langle W_i^\eta(t)\eta_i(t')\rangle = D_i^m \chi_i^\eta + \sigma^2 \int_{-\infty}^{t'} ds C_i^\xi(s,t)\chi_i^\eta(t',s)\,. \tag{S.59}$$

With the choice $t' > t$ we arrive to $\mu_i^\eta(t) = -\gamma_i + \sigma^2 \int_{-\infty}^{t'} ds C_i^\xi(s,t)\chi_i^\eta(t,s) + D_i^m/2$. We have only to evaluate equal time cross correlations, which are given by

$$\begin{aligned}\partial_t C_i^{\xi\eta} &= g_i C_i^\eta - d_i C_i^{\xi\eta} - \mu_i^\xi(t) C_i^{\xi\eta}\,,\\ \partial_t C_i^{\xi\eta} &= -\gamma_i C_i^{\xi\eta} - \mu_i^\eta(t) C_i^{\xi\eta}\,,\end{aligned} \tag{S.60}$$

and subtracting the two equations we arrive to

$$C_i^{\xi\eta} = \frac{g_i C_i^\eta}{d_i - \gamma_i + \mu^\xi(t) - \mu_i^\eta(t)}\,. \tag{S.61}$$

It has to be noticed that $\mu^\xi(t)$ depends on the equal cross-correlation,

$$C_i^{\xi\eta} = \frac{g_i C_i^\eta}{g_i C_i^{\xi\eta} + \frac{D_i^n}{2} - \sigma^2 \int_{-\infty}^{t'} ds C_i^\xi(s,t)\chi_i^\eta(t,s) - \frac{D_i^m}{2}}\,. \tag{S.62}$$

We close the equation for the spherical constraints upon solving the equation for the response of mRNA fluctuations,

$$\partial_t \chi_i^\eta = -(\gamma_i + \mu_i^\eta(t))\chi_i^\eta + \delta(t,t')\,. \tag{S.63}$$

Once we found the equation for the spherical constraint the equation for the correlations are compactly given by

$$\begin{aligned}\partial_t C_i^\xi &= g_i(C^{\xi\eta} - C_i^{\xi\eta} C^\xi(t,t')) - \frac{D_i^n}{2} C_i^\xi + D_i^n \chi^\xi\,,\\ \partial_t C_i^\eta &= -\frac{D_i^m}{2} C^\eta + D_i^m \chi_i^\eta + \sigma^2(1 - C_i^\eta)\int_{-\infty}^{t'} ds C_i^\xi(s,t)\chi_i^\eta(t,s)\,,\end{aligned} \tag{S.64}$$

In the following, we drop the index $i$ and take delta distributions of the parameters. For long enough times, we look for solutions such that the fluctuation-dissipation theorem holds, and so the last equation is ($t' > t, t' - t = \tau$),

$$\begin{aligned}\partial_t C^\xi(\tau) =& g\left(C^{\xi\eta}(\tau) - C^{\xi\eta}(0)C^\xi(\tau)\right) - \frac{D^n}{2} C^\xi(\tau)\,,\\ \partial_t C^\eta(\tau) =& -\frac{D^m}{2} C^\eta(\tau)\\ &- \frac{2\sigma^2}{D^m}(1 - C^\eta(\tau))\int_0^\tau du C^\xi(\tau - u)\partial_u C^\eta(u)\,.\end{aligned} \tag{S.65}$$

The last integral can be approximated for $\tau \gg 1$ ,

$$\begin{aligned}\partial_\tau C^\xi(\tau) &= g\left(C^{\xi\eta}(\tau) - C^{\xi\eta}(0)C^\xi(\tau)\right) - \frac{D^n}{2}C^\xi(\tau)\,,\\ \partial_\tau C^\eta(\tau) &= -\frac{D^m}{2}C^\eta(\tau) + \frac{2\sigma^2}{D^m}(1-C^\eta(\tau))^2 C^\xi(\tau)\,.\end{aligned} \tag{S.66}$$

As we are looking at a relaxation problem we expect $\partial_\tau C^{\xi\setminus\eta}(\tau) \leq 0$ and find $C^{\xi\eta}(0) \approx 1$ (see (S.62)). In this regime the following inequalities must hold (upon substituting the solution of $C^{\xi\eta}$):

$$\begin{aligned}&\frac{4\sigma^2}{(D^m)^2}C^\xi(\tau)(1-C^\eta(\tau)) > \frac{2g+D^n-D^m}{D^m}\\ &C^\xi(\tau)(D^n+2g)\Big[D^m - D^n - 2g\\ &\quad + \frac{2\sigma^2}{D^m}C^\xi(\tau)\,(1-C^\eta(\tau))\Big] + 4C^\eta(\tau)g^2 \leq 0\end{aligned} \tag{S.67}$$

As $D^m \gg D^n$ and $D^m \gg g$ the second inequality is always satisfied. Instead, the first inequality simplifies to,

$$\frac{4\sigma^2}{(D^m)^2}C^\xi(\tau)(1-C^\eta(\tau))^2 \leq C^\eta(\tau)\,. \tag{S.68}$$

In particular, for $\tau = 0$, $C^\xi(0) = C^\eta(0) = 1$ as a consequence of the spherical constraint, which implies that, in the same limits, the previous inequality is not satisfied when

$$\sigma_c = D^m/2\,. \tag{S.69}$$

Equation (S.69) defines a dynamical transition in the autocorrelation functions. In order to compare these results with the phase diagram (Fig. 1) we have to divide $\sigma$ by $D^m$ and the critical line is defined by $\sigma_c = 1/2$.
