## Supplementary Figures and Theory for "Coordinated gene expression variability encodes the regulatory state of cells": Supplementary_Figures.pdf

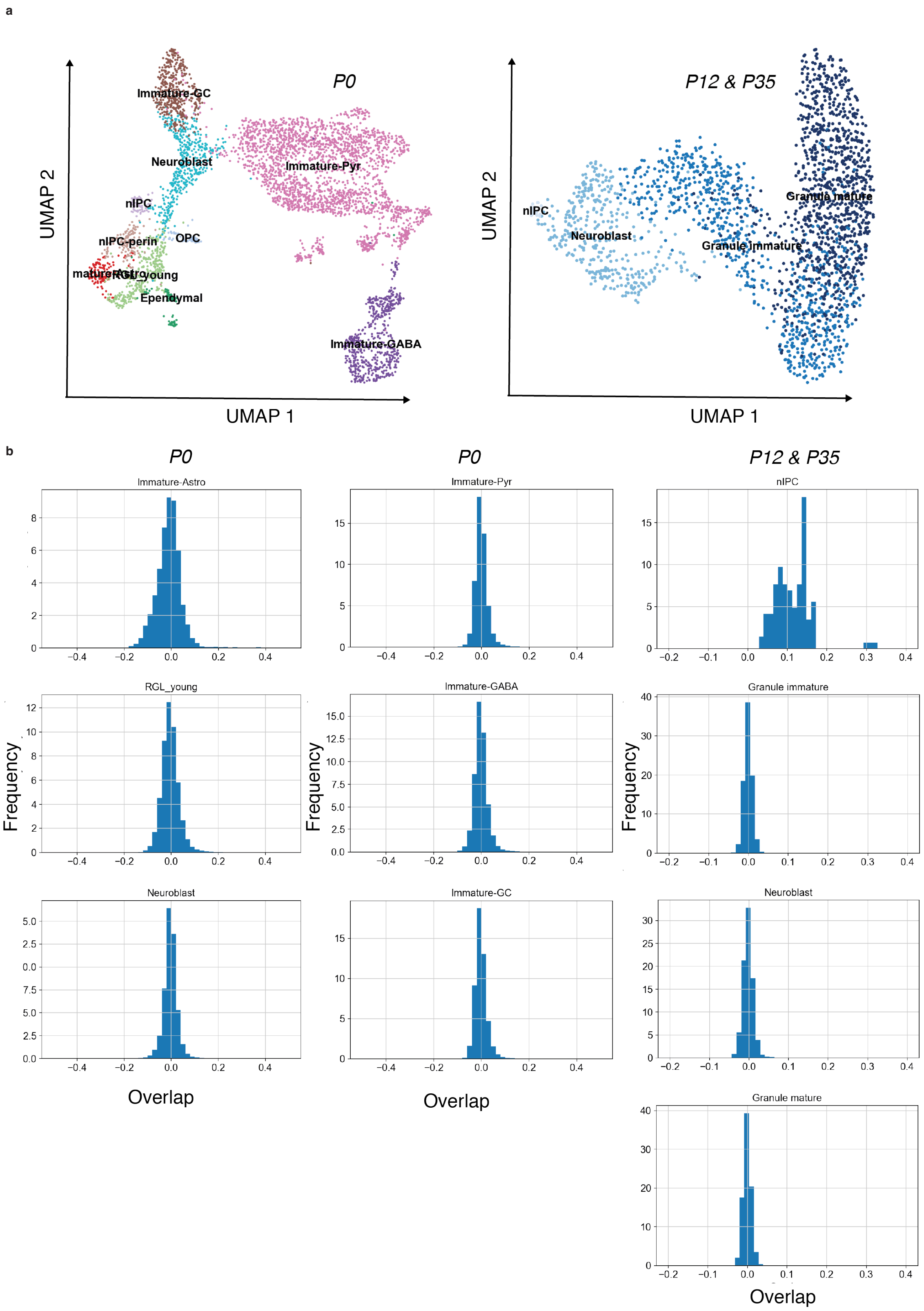


Figure S1: (**a**) Clusters with the respective annotation from the data in [1] of developing mouse brain. (**b**) Overlap distribution for each of the clusters in **a**.

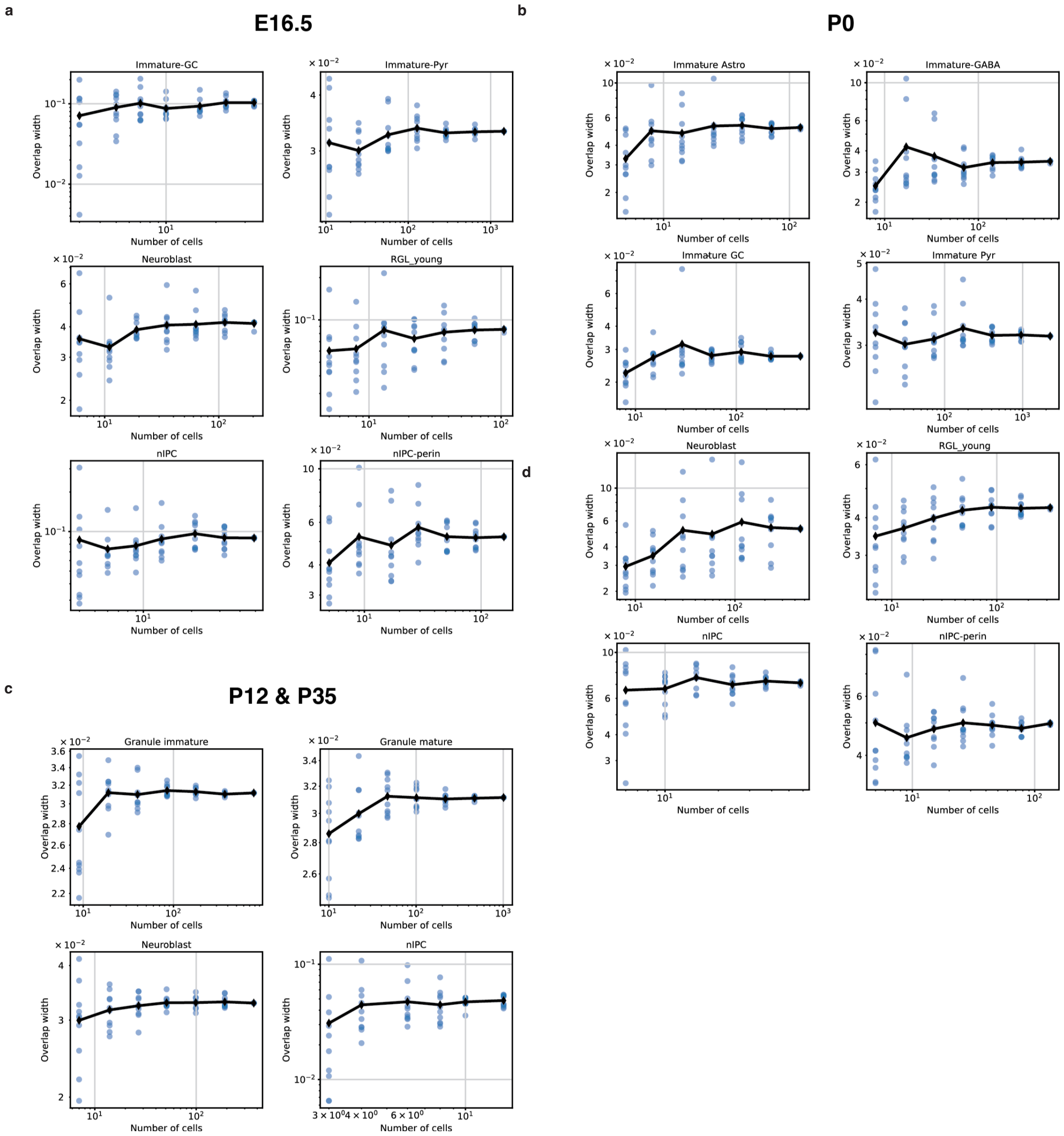


Figure S2: Dependence of overlap distribution width (standard deviation) on the number of cells in a cluster. For each of the clusters from the three data sets (**a**): E16.5, (**b**): P0, (**c**): P12 & P35, we subsampled cells by randomly choosing cells and computing their KNN neighborhoods. Each KNN neighborhood is identified as a new steady state and we compute overlaps for each such configuration. The average overlap width (black line) does not depend appreciably on the number of cells.

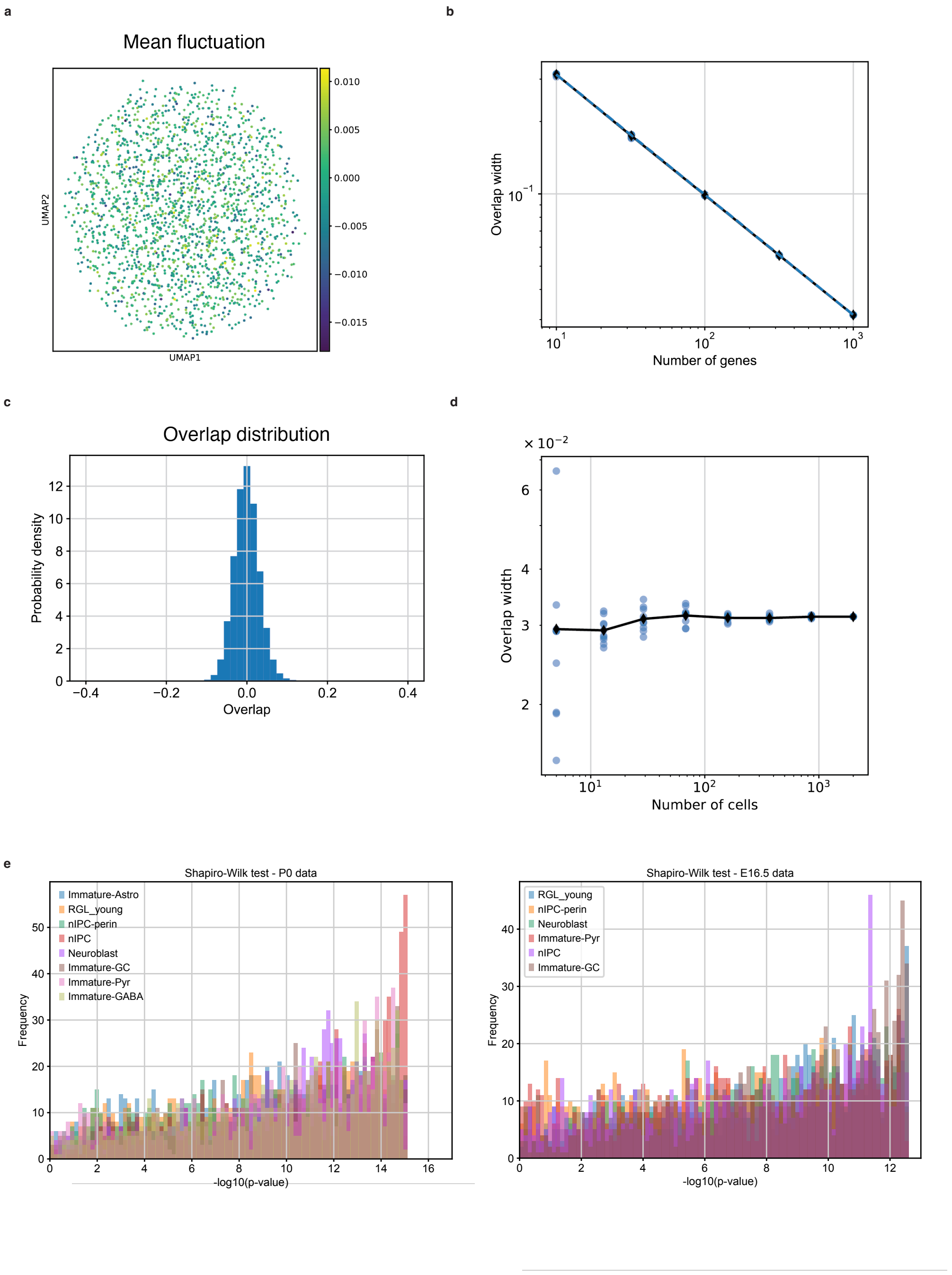


Figure S3: Simulated uncorrelated cells do not show correalated structure. (**a**) We simulated cells using *splatter* and confirmed that the cells are uncorrelated because the mean fluctuations (averaged over all genes) show no pattern. (**b**) The overlap width scales as $1/\sqrt{N}$ as expected from the central limit theorem. (**c**) The overlap distribution is approximately normal and is unimodal. (**d**) Subsampling cells using KNN neighbourhoods confirms that the overlap width does not depend on number of cells in a cluster. (**e**) Shapiro-Wilk test for normality of fluctuations on the representative clusters in Fig. 2.

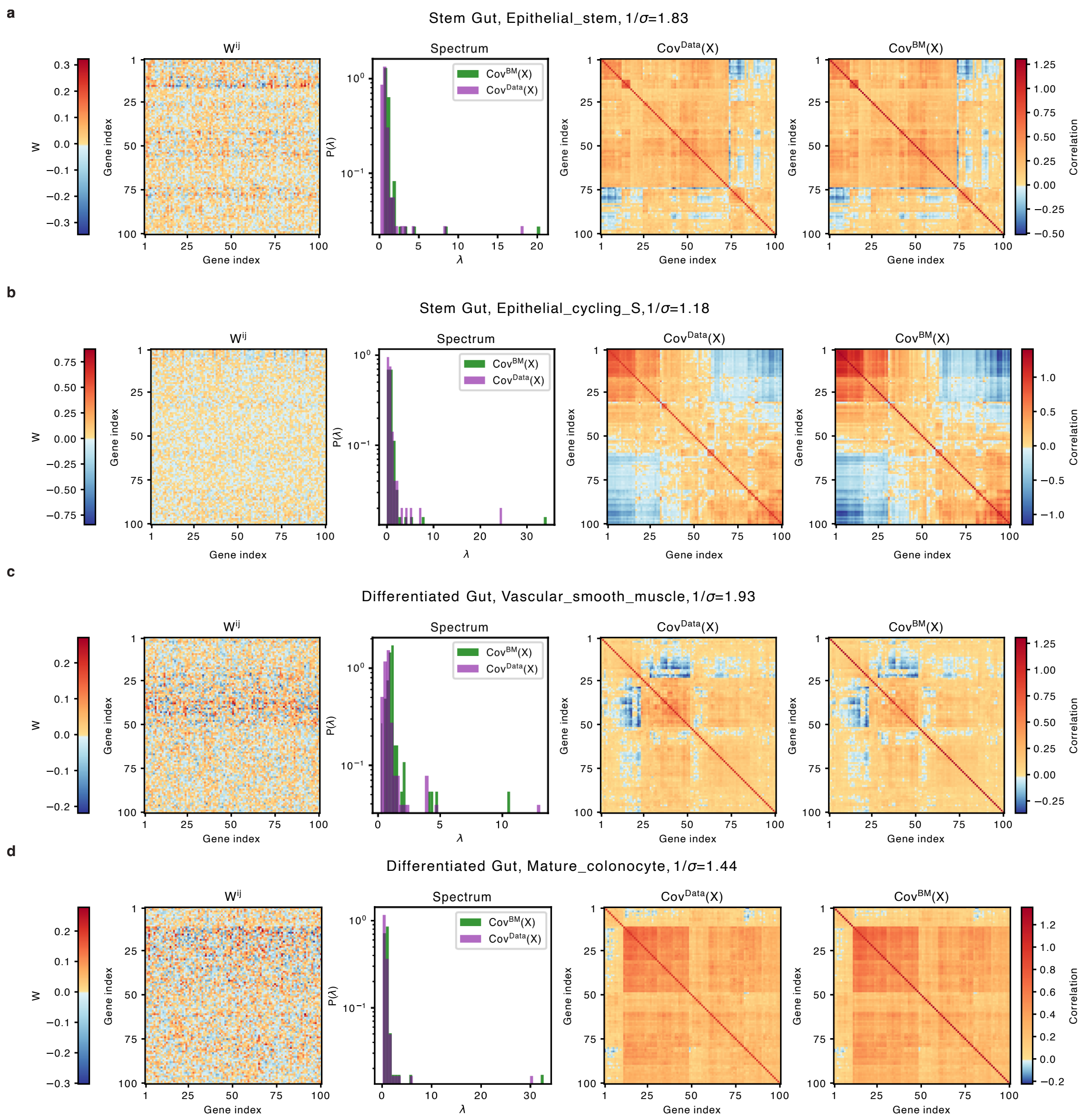


Figure S4: From left to right: Interaction matrix obtained from the RBM, Spectrum of the eigenvalue for the experimental and RBM derived covariance matrices, experimental covariance matrices and . The cell clusters are shown in Fig. 3 and defined in Methods §1.2. The inference for all the remaining annotated clusters are provided in [2].

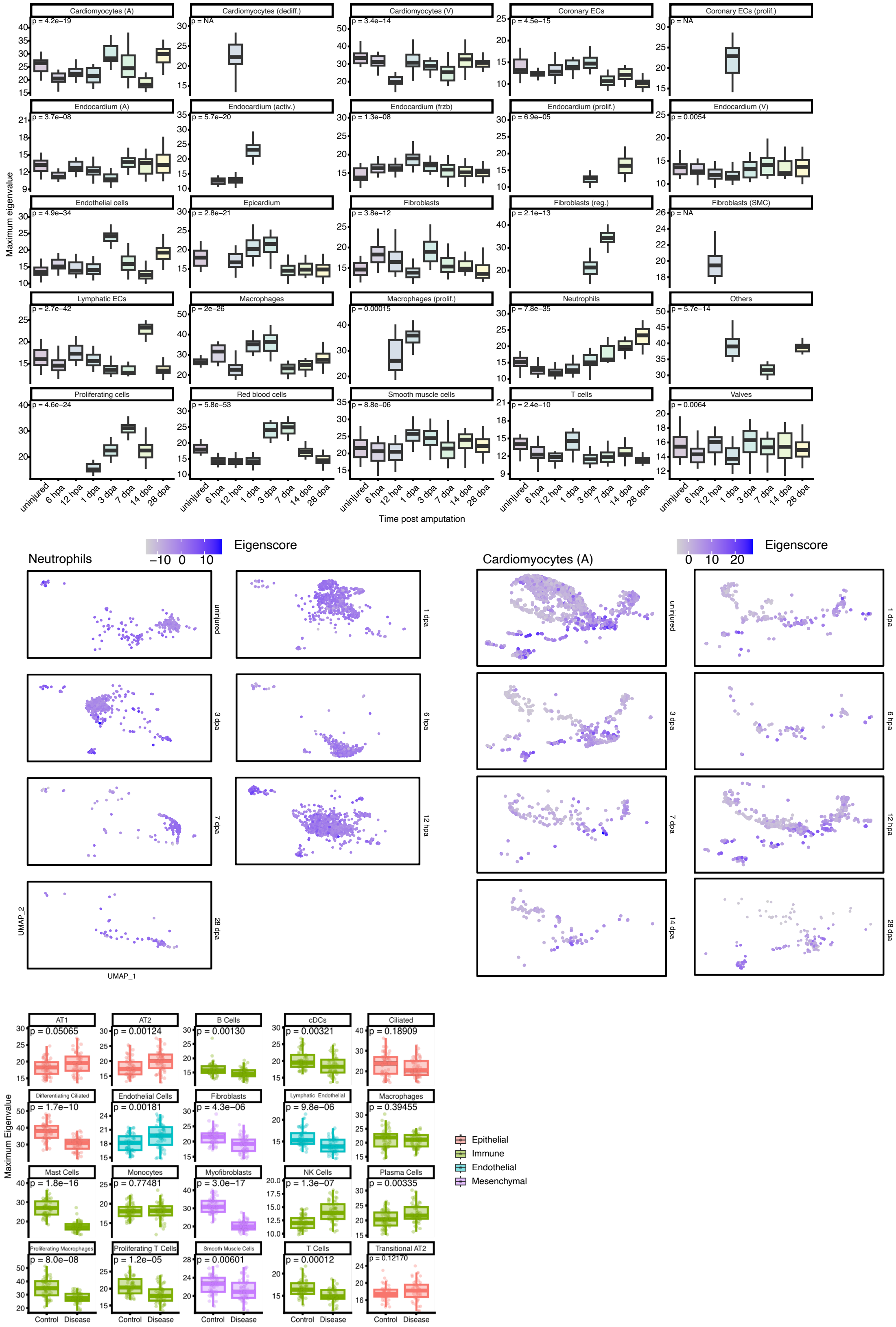


Figure S5: (**a**) Maximum eigenvalue computed with bootstrapping of clusters in Fig. 4a,b as explained in Methods§1.3. p-values are computed with the ANOVA test in each cluster between days. (**b**) Projection of the eigenvector associated to the maximum eigenvalue for each cell in annotated Neutrophilis and Cardiomyocytes (A) clusters. There is no clear sub-population dominating the score. (**c**) Maximum eigenvalue computed with bootstrapping of clusters in Fig. 4c,d as explained in Methods§1.3. p-values are computed with the Mann U-Whiteny test in each cluster between the two conditions (Control and Disease).
